# Single molecule MATAC-seq reveals key determinants of DNA replication origin efficiency

**DOI:** 10.1101/2023.03.14.532513

**Authors:** Anna Chanou, Matthias Weiβ, Karoline Holler, Tobias Straub, Jana Krietsch, Andrea Sanchi, Henning Ummethum, Clare S. K. Lee, Elisabeth Kruse, Manuel Trauner, Marcel Werner, Maxime Lalonde, Massimo Lopes, Antonio Scialdone, Stephan Hamperl

**Affiliations:** Institute of Epigenetics and Stem Cells, Helmholtz Zentrum München, Munich, Germany; Core Facility Bioinformatics, Biomedizinisches Centrum (BMC) LMU Munich, Germany; Institute of Functional Epigenetics, Helmholtz Zentrum München, Neuherberg, Germany; Institute of Computational Biology, Helmholtz Zentrum München, Neuherberg, Germany; Institute of Molecular Cancer Research, University of Zurich, Zurich, Switzerland

## Abstract

Stochastic origin activation gives rise to significant cell-to-cell variability in the pattern of genome replication. The molecular basis for heterogeneity in efficiency and timing of individual origins is a long-standing question. Here, we developed **M**ethylation **A**ccessibility of **TA**rgeted **C**hromatin domain Sequencing (MATAC-Seq) to determine single-molecule chromatin accessibility of specific genomic loci after targeted purification in their native chromatin context. Applying MATAC-Seq to selected early-efficient (EE) and late-inefficient (LI) budding yeast replication origins revealed large heterogeneity of chromatin states. Disruption of INO80 or ISW2 chromatin remodeling complexes leads to changes at individual nucleosomal positions that correlate with changes in their replication efficiency. We found a chromatin state with an optimal 100-115bp nucleosome-free region in combination with surrounding well-positioned nucleosomes and open +2 linker region is a strong predictor for efficient origin activation. Thus, MATAC-Seq identifies the large spectrum of alternative chromatin states that co-exist on a given locus previously masked in population-based experiments and provides a mechanistic basis for origin activation heterogeneity during DNA replication of eukaryotic cells. Consequently, our single-molecule assay for chromatin accessibility will be ideal to define single-molecule heterogeneity across many fundamental biological processes such as transcription, replication, or DNA repair *in vitro* and *ex vivo*.

## Introduction

In eukaryotic genomes, multiple start sites of DNA replication named origins are distributed across each of the linear chromosomes. The assembly of replication forks and the bidirectional initiation of DNA synthesis at these sites is known as origin firing. The budding yeast *Saccharomyces cerevisiae* has been a particularly useful model organism to study eukaryotic replication, owing to the presence of short (∼200bp) Autonomous Replication Sequences (ARS) that support the propagation and maintenance of plasmids and therefore defined chromosomal replication origins (9, 10). At the core of every yeast replication origin is a replicator sequence containing a conserved AT-rich 11bp stretch known as the ARS consensus sequence (ACS), which is essential for binding of the initiator complex known as the origin recognition complex (ORC). This six-subunit complex (Orc1-6) is highly conserved among eukaryotes and works in concert with Cdc6 and Cdt1 to direct the loading of the replicative helicase Mcm2–7 onto DNA to form the pre-replicative complex (pre-RC). The assembly of the pre-RC in G1 serves to “license” an excess of origins for activation in the subsequent S-phase. Cyclin- and Dbf4-dependent kinase activities during S-phase result in the recruitment of additional proteins at a fraction of loaded pre-RCs to form the preinitiation complex, to unwind the DNA, and produce bidirectional replication forks (11).

Similar to all other eukaryotic genomes, yeast replication origins can be classified according to two main intrinsic features, replication timing and efficiency. Replication timing denotes the timepoint when an origin fires and can occur at a continuum from early to late S-phase (12, 13). Replication efficiency describes the probability of activation for each replication origin (i.e. the fraction of cells in which the origin fires) and is defined as a combination of an origin’s firing probability and its proximity to other origins that fire earlier and can therefore passively replicate the origin. Origin efficiencies display a wide range from inefficient (≤ 10%) to highly efficient (≥ 90%) (7). The difference in replication timing and efficiency of individual origins scattered across the chromosomes defines the replication timing program that shows a highly reproducible order of chromosomal replication at the population level. However, at the single cell level, origin firing is an intrinsically stochastic event with no two chromosomes exhibiting the same replication pattern in an isogenic cell population (14, 15). The observed stochasticity of origin activation eventually leads to further cell-to-cell variability (1–8). This apparent contradiction can be explained by the fact that the population experiments average the behavior of individual origins over large numbers of cells, obscuring stochastic effects. Thus, the molecular rules based on which individual cells choose to activate a specific, highly individualized subset of replication origins remains an outstanding fundamental question (1, 16).

A plausible mechanism by which firing probability could be regulated is the local chromatin structure. Yeast ARS sequence is AT-rich which is inhibitory to nucleosome assembly and forms a nucleosome-free region (NFR). An NFR at an ARS provides an accessible environment for ORC binding and replication initiation(17, 18). Importantly, ORC-binding at functional ARS improves nucleosomal positioning of the -1 and +1 nucleosomes surrounding the ARS. Disruption of this ORC-directed nucleosome phasing by mutating the ORC binding site or moving one nucleosome further from the ARS interferes with the efficiency of origin firing (18, 19). Genome-wide nucleosome positioning maps in G1-phase revealed many different clusters of distinct nucleosome occupancy patterns around annotated origins (20), suggesting large heterogeneity of different chromatin states. A later study showed that early-origins tend to show higher nucleosome occupancy at the +1 and -1 position around the ARS sites compared to late origins (21), suggesting that the interaction of ORC with origin DNA and neighboring nucleosomes is critical for efficient and timely replication. This is also consistent with recent *in vitro* reconstitution experiments (22) that show stable ORC-nucleosome complexes that can efficiently load the MCM2-7 double hexamer onto adjacent nucleosome-free DNA. Neighboring genomic features, most notably transcription start sites (TSS) and gene ends, were also shown to influence the nucleosome positioning and thus the replication identity of an origin (20). Together, these data suggest a model of nucleosome positioning at replication origins in which the underlying ARS sequence occludes nucleosomes to permit binding of ORC, which then likely in concert with nucleosome modifiers or chromatin remodeling enzymes (CREs) position nucleosomes at origins in a favorable way to promote replication origin function.

Because CREs are major architects of chromatin structure(23–32), we hypothesized that comparing chromatin accessibilities in wildtype and CRE mutant strains at selected replication origins will allow us to extract the key differences in chromatin states that are important for the functional activation of DNA replication origins. However, critical testing of this hypothesis necessitates the analysis of origin chromatin structure at the level of single cells or single DNA molecules. To determine the precise nucleosome configuration(s) restricting or allowing replication initiation, we developed **M**ethylation **A**ccessibility of **TA**rgeted **C**hromatin domain **Seq**uencing (**MATAC-Seq**) to define single-molecule chromatin accessibility maps of specific genomic loci after targeted purification in their native chromatin context. Applying MATAC-Seq to selected early-efficient (EE) and late-inefficient (LI) budding yeast replication origins revealed large cell-to-cell heterogeneity in their chromatin states. Disruption of INO80 or ISW2 chromatin remodeling complexes leads to changes at individual nucleosomal positions that correlate with changes in their replication efficiency. Thus, MATAC-Seq identifies the large spectrum of alternative nucleosome occupancy states that co-exist on a given locus. In replication origins, this allowed us to extract key chromatin features of stochastic origin activation previously masked in common population-based techniques and provides a mechanistic basis for origin activation heterogeneity during DNA replication of eukaryotic cells.

## Results

### ISW2 and INO80 chromatin remodelers affect replication efficiency of selected EE and LI replication origins

To test this hypothesis, we focused on four selected replication origins on yeast chromosome III, which display previously characterized major differences in their replication timing and efficiency properties(33, 34) (**Fig. 1A**). We analyzed their replication timing properties in wildtype and CRE mutant strains deleted for subunits of the ISW2 (isw2Δ) and INO80 (ies6Δ) remodeling complexes using quantitative PCR (qPCR)-based DNA copy number assay (**Fig. 1B**). This analysis of the early-efficient (EE) ARS305 and ARS315 and late-inefficient (LI) ARS313 and ARS316 origins over a late replicating region on chromosome IV(35) showed the expected early or late replication profiles in wildtype cells (**Fig. 1C-F**). However, both CRE mutant strains displayed lower replication efficiency at both EE origins. For example, ARS305 copy number dropped by 19% in isw2Δ and 18% in ies6Δ 32 min after release (**Fig. 1C**). Similarly, ARS315 copy number dropped by 23% in isw2Δ and 18% in ies6Δ 40 min after release (**Fig. 1D**). Interestingly, the LI origin ARS313 displayed 20% higher replication efficiency in ies6Δ at 32 min, whereas no significant effect was observed for the isw2Δ mutant **(Fig. 1E**). In contrast, the replication profile of ARS316 was unaffected in both CRE mutant strains (**Fig. 1F**). Growth curves and FACS profiles showed that these changes in replication efficiency were not caused by growth defects (**Fig. S1A**) or differences in global arrest and release kinetics between wildtype and mutant strains (**Fig. S1B**). These results suggested that ISW2 and INO80 CREs impact the replication efficiency of individual origins but do not globally change the replication program and S-phase progression.

**Figure 1.**
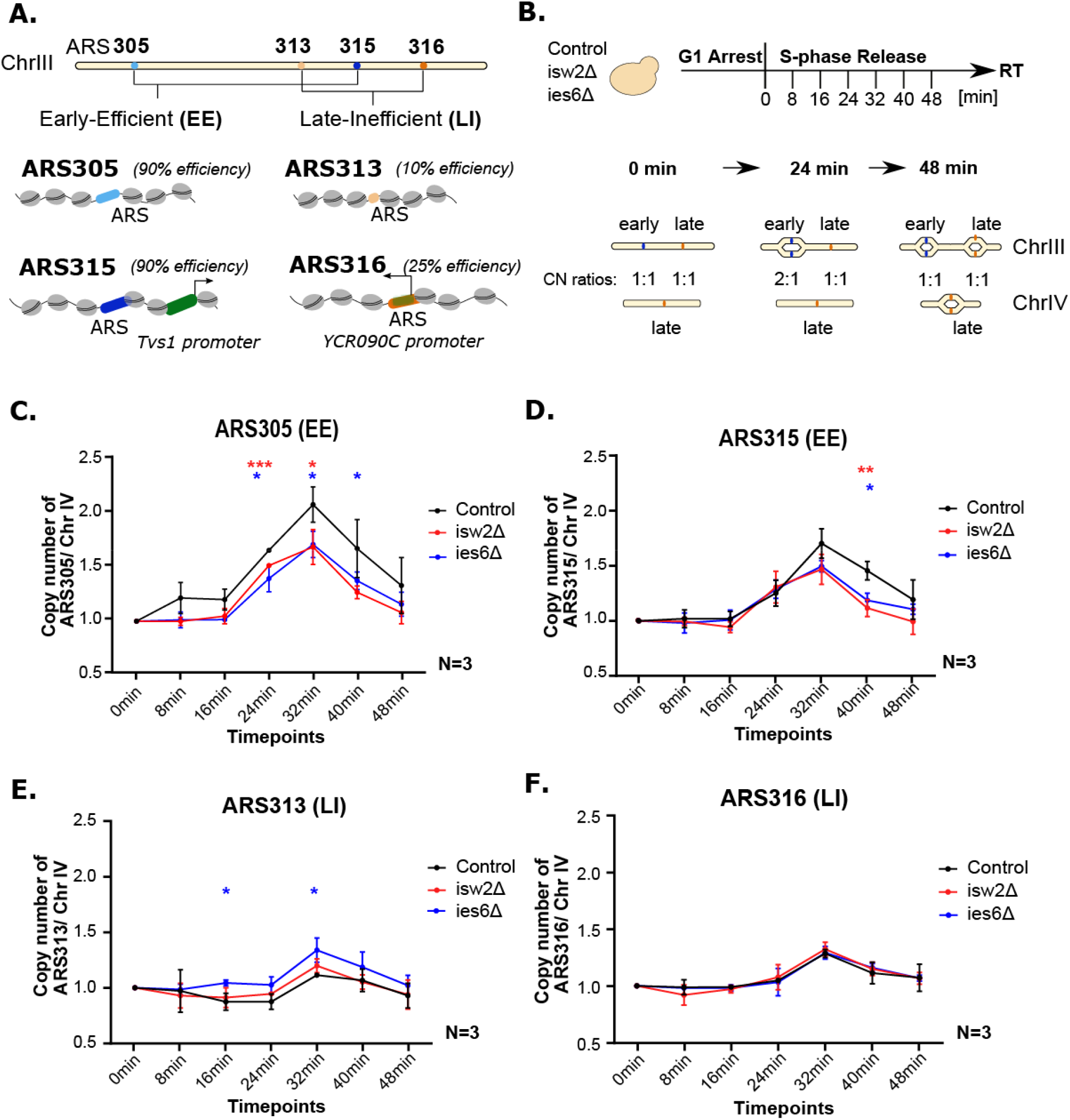
**A.** Schematic representation of Early-Efficient (EE) and Late-Inefficient (LI) replication origins on Chromosome III of *Saccharomyces cerevisiae*. The two EE origins (ARS305/ARS315 – ARS blue) show 90% efficiency and two LI origins (ARS313/ARS316 – ARS orange) show 10% and 25% efficiency, respectively. Distinct chromatin features are indicated around the ARS loci such as nucleosomes (grey) and gene promoters (green). **B.** Experimental outline for measuring replication timing of each origin (Control, isw2Δ, ies6Δ). After 2 h arrest, the cells were released into S-phase and samples were collected every 8 min for FACS (**Fig. S1B**) and qPCR. A late replicating region of ChrIV is used as reference. The ratio between the origin of interest and the reference region shows the replication efficiency. **C.-F.** The replication timing plots show the average copy number ratios of ARS305, ARS315, ARS313, ARS316 to the late replicating region of ChrIV with standard deviation from n = 3 biological replicates (*, **, *** indicates statistical significance P < 0.05, or P < 0.01, or P < 0.001, respectively, by unpaired t-test).

### Bulk chromatin accessibility analysis shows large differences between origins and ies6Δ mutant cells

As a first approach to measure bulk nucleosome occupancy at our selected replication origins, we applied classical restriction enzyme (RE) accessibility assays (36–38) (**Fig. 2A**) at individual positions around the ARS expected to be protected by NS-2, NS-1, NS+1 or NS+2 nucleosomes around the ARS (**Fig. 2B**). For this study, we define the +1 nucleosome of an origin as the nucleosome that is closest to the annotated ACS within the ARS where the initial binding of ORC occurs. Nuclei from logarithmically growing wildtype strain were digested at different RE concentrations to establish saturation and analyzed by Southern blot analysis with specific probes directed against the selected EE and LI origins. In the wildtype strain, we observed similar percentages of digested molecules (∼30-40%) at the NS+1 and NS+2 restriction sites among all 4 origins (**Fig. 2C-E**), indicating that around 30-40% of all molecules were not protected by nucleosomes at these two positions. Interestingly, we observed major differences in RE accessibility between origins at the NS-2 and NS-1 positions. For example, the NS-2 HpaI site only showed 20% digestion at ARS305, whereas more than 60% of molecules were digested by HpaI in the same relative NS-2 position of ARS316 (**Fig. 2C**). We also note that the ARS305 NS-1 position showed strong protection (∼10% digestion), whereas more than 50% of the molecules were accessible at this site in ARS316 (**Fig. 2C**), consistent with a strongly positioned NS-1 nucleosome at the EE origin ARS305 (21).

**Figure 2.**
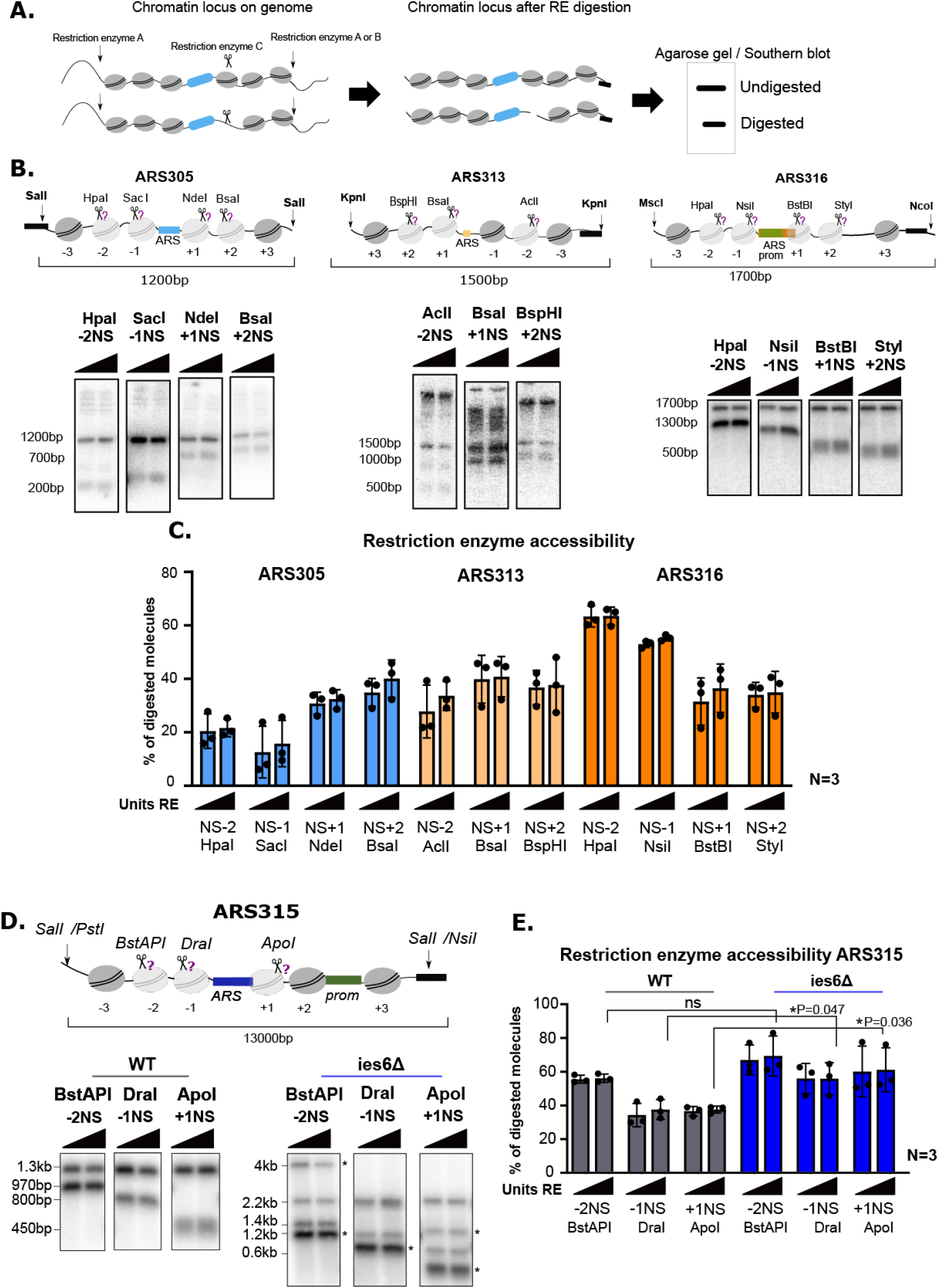
**A.** Cartoon showing the principal of Restriction Enzyme Accessibility (REA) assay **B.** Restriction endonuclease accessibilities in chromosomal ARS305, ARS313 and ARS316 locus. Nuclei from yeast strains Y65 (ARS305), Y94 (ARS313) and Y69 (ARS316) were isolated and digested with increasing amounts (10U, 50U or 100U) of the indicated restriction enzymes (triangle on top of each pair of panels / scissors). DNA was isolated, digested with SalI in ARS305, KpnI in ARS313 and MscI/NcoI in ARS316 and subjected to indirect end-labeling Southern blot analysis with the radioactively labeled probe. **C.** The histogram shows the results of Southern blot quantification as a percentage of digested chromatin locus. **D.** Restriction endonuclease accessibilities (REAs) in chromosomal ARS315 locus. Nuclei from yeast strains Y91 (control-ARS315), and Y130 (ies6Δ-ARS315) were isolated and digested with increasing amounts (10U, 50U and 100U) of the indicated restriction enzymes (scissors). DNA was isolated, digested with SalI and subjected to indirect end-labeling Southern blot analysis with the radioactively labeled probe. Top: schematic representation of the ARS315 locus with restriction sites used to probe chromatin structure (scissors) and secondary restriction sites to isolate the locus. Asterisks in the mutant strain indicate restriction bands derived from a spike-in naked plasmid control to verify complete digestion of the enzymes (see also **Fig. S2**). **E.** The histogram shows the results of Southern blot quantification as a percentage of digested chromatin locus. Average and standard deviations are from n= 3 biological replicates (*, **, *** indicates statistical significance P < 0.05, or P < 0.01, or P < 0.001, respectively, by unpaired t-test).

We next asked whether knockout of IES6 led to changes in absolute nucleosome occupancy at selected sites around ARS315 (**Fig 2D-E**). In the ies6Δ samples, we also included in the RE accessibility assay a naked spike-in plasmid DNA containing the same ARS315 origin DNA sequence and observed 100% digestion of the naked DNA into the expected DNA fragments (**Fig. 2D, DNA bands marked with asterisks and Fig. S2**), confirming the reactivity of the used REs and that the fraction of remaining undigested molecules from the nuclei was caused by the chromatin state of ARS315. We observed no difference in accessibility between wildtype and ies6Δ strain at the NS-2 BstAPI restriction site located 222 bp upstream of the ARS **(Fig. 2E**). However, the DraI and ApoI sites flanking the ARS in NS-1 and NS+1 positions showed an 18% and 24% increase in accessibility, respectively (**Fig. 2E**), suggesting that upon loss of INO80, the typically well positioned nucleosomes in close vicinity to the ARS are evicted or repositioned, thereby making this site more accessible in the mutant strain. Together, these bulk analyses suggested an inherent level of heterogeneity in nucleosomal occupancy at individual positions and that INO80 can affect chromatin accessibility at ARS315.

### Functional replication origins are purified with high yield and purity in their native chromatin context

To obtain a more comprehensive readout of chromatin accessibility and nucleosome occupancy changes, we sought to investigate potential local and global changes of the chromatin landscape of the targeted replication origins between CRE mutant and wildtype strains. Ideally, the method of choice should allow for quantitative profiling of the accessibility of individual chromatin fibers at the single-molecule level with high resolution and coverage, thereby revealing the heterogeneity of multiple chromatin states that may co-exist in a cell population. For this, we developed **M**ethylation **A**ccessibility of **TA**rgeted **C**hromatin domain Sequencing (**MATAC-Seq**), a single-molecule approach to profile the chromatin structure of a targeted locus of interest at near basepair resolution. MATAC-Seq is built on the conceptual foundations of NOMe-Seq and SMAC-Seq (39–43) that rely on the preferential modification of accessible DNA by DNA methyltransferases combined with Nanopore sequencing for direct readout of the methylated DNA bases. In MATAC-Seq, we additionally take advantage of our previously established single-copy locus chromatin purification approach(44–46). This strategy allows strong enrichment for our targeted replication origins prior to methylation footprinting and Nanopore sequencing analysis, thereby overcoming current limitations of DNA sequencing coverage of similar genome-wide approaches (42).

Briefly, site-specific recombination allows excision of native chromatin rings encompassing a ∼ 1kb genomic region centered around the single-copy replication origins from its endogenous chromosomal location in G1-phase arrested cells (**Fig. 3A-B**). Insertion of RS/LEXA sites had no impact on growth and generation timing in all recombination-competent strains compared to the control strain (**Fig. S3A**). We monitored the recombination kinetics and efficiency under these conditions in a time-course experiment by Southern blotting of extracted genomic DNA in a negative Control strain (no RS/LexA sites) and the recombination-competent ARS315 and ARS313 strains. We observed near complete recombination of the targeted locus within ∼60-90 min after recombinase induction (**Fig. 3C-D and Fig. S3B-D**). Cells were lysed and the whole cell extract subjected to a single-step affinity purification protocol using the LexA-TAP protein as an affinity handle (**Fig. 3A**). We took DNA samples from the resulting Input (IN), Flowthrough (FT), Beads (B) and Elution (E) fractions and quantified the amount of ARS315 DNA by qPCR (**Fig. 3E**). After Tobacco Etch Virus (TEV) protease-mediated cleavage of LexA-TAP on the beads, ∼ 15% of all chromatin rings relative to the input were recovered in the final elution sample, whereas no enrichment of ARS315 was observed in the control strain. Indeed, the purified eluate fraction showed a strong enrichment of ARS315 over an unrelated control locus and we estimate that ∼70-80% of all DNA molecules in these samples were derived from our targeted loci (**Fig. 3F**).

**Figure 3.**
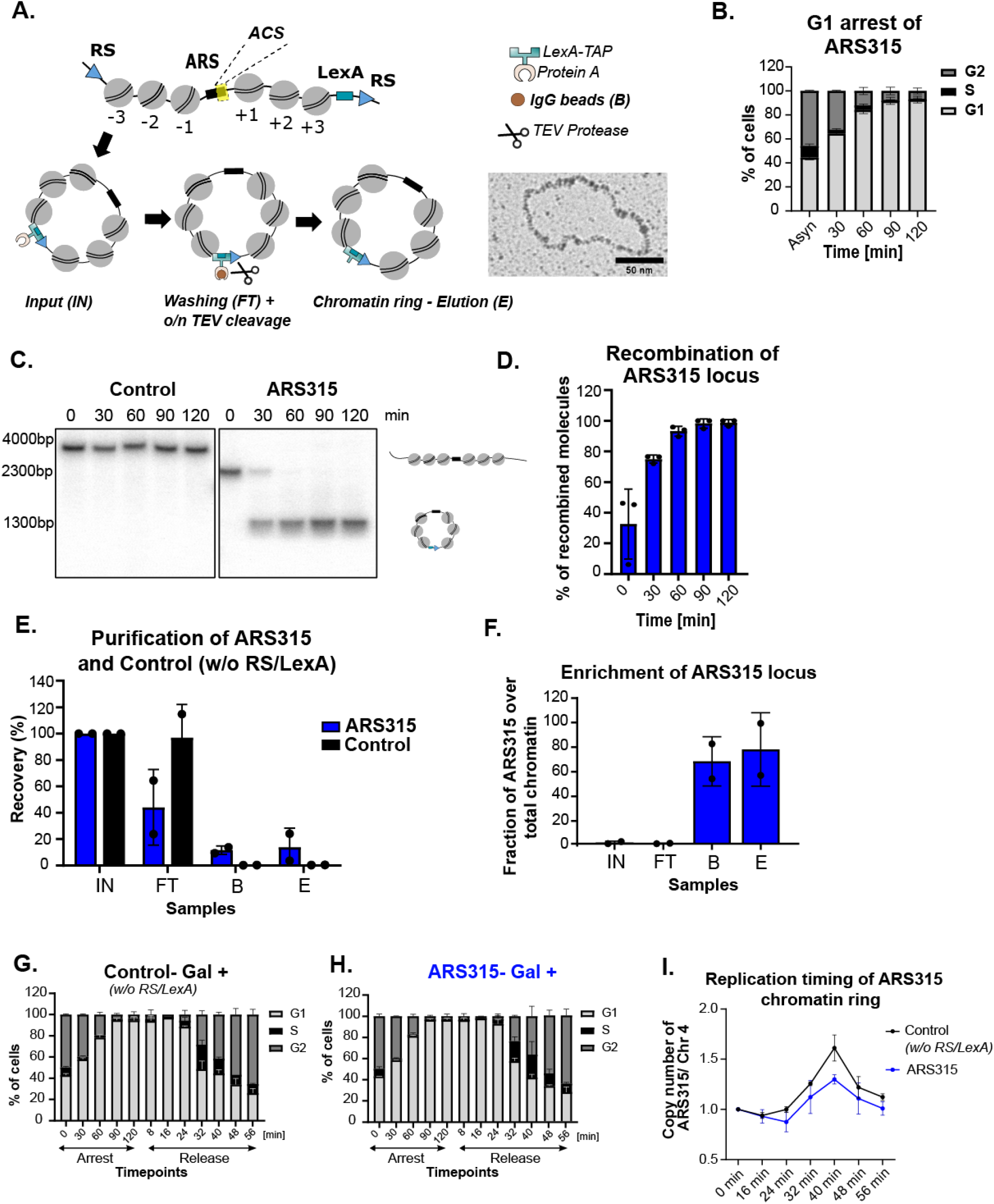
**A.** Schematic representation of LexA affinity purification. Recombination sites (RS) were integrated after the +/- 3 nucleosomes around the ARS locus of single copy origins and after the +/-2 nucleosomes of the multi-copy rARS origin. By convention, the nucleosomes located closer or further away to the ACS site of the ARS was termed as (+) or (-), respectively. Electron micrograph shows individual ribosomal ARS molecule after native spreading. **B.** Cell arrest in G1-phase. The strain Y91-ARS315 was grown in YPR medium and arrested with 50 ng/ml α-factor. Samples for FACS analysis were taken at the indicated timepoints and strained by Sytox red to monitor the distribution of G1, S and G2-phase in both profiles. **C.** Recombination kinetics of ARS315 locus. The strains Y91-ARS315 and Y66-control (w/o RS or LexA sites) were grown in YPR medium to logarithmic phase and arrested in G1 phase by addition of α-factor (50ng/ml) and recombination was induced by the addition of 2% galactose. Samples were taken at indicated timepoints. DNA was isolated and linearized by BsrGI and subjected to Southern blot analysis. The positions of unrecombined and recombined molecules are shown on the right. **D.** The histogram shows the results of Southern blot as percentage of recombined chromatin locus. **E.** LexA affinity purification was performed for strain Y91, where the single-copy locus is targeted by RS, and strain Y66 (control), lacking RS/LexA recognition sites but containing the LexA expression cassette, therefore no recombination or purified chromatin is expected. DNA samples were taken (0.1% Input (IN), Flowthrough (FT), Beads (B), Eluate (E) from n=2 biological replicates and analyzed by qPCR. The enrichment of an unrelated region (PDC1) was analyzed side-by-side with the regions of interest. **F.** Enrichment of ARS315 locus in the final elute. Given that the size of yeast genome is 12 kb, and the length of a chromatin ring is ∼1 kb, the fold enrichment ratio of the specific origin to PDC1 was used to define the enrichment of an origin in the DNA samples. **G-H.** Cell cycle progression of control (Y66) (w/o RS/LexA sites) and ARS315 (Y91-with RS/LexA sites) strains after addition of 2% galactose. The cells were arrested by 50 ng/ml α-factor and released into replication by 125U Pronase. The FACS samples and qPCR samples were taken side-by-side at the indicated timepoints. **I.** The replication timing plots show the average copy number ratios after addition of 2% galactose of ARS315 locus, either in the Control (Y66) strain containing no RS/LexA sites for recombination or in the ARS315 (Y91) containing RS/LexA sites, to the late replicating region of ChrIV with standard deviation from n = 2 biological replicates. The cells have been arrested for 2 h by α-factor and released to S-phase by 125U Pronase.

Importantly, we verified that our genetic manipulations and the necessary process of recombination did not interfere with functionality of the origin. To this end, control cells without RS/LexA and ARS315 cells were arrested in G1-phase with alpha factor with addition of Galactose (Gal +) to induce recombination in the ARS315 strain and then released synchronously into S-phase. FACS analysis showed highly similar arrest and release kinetics between the two strains (**Fig. 3G-H**). We then performed DNA copy number analysis at different timepoints after release by qPCR comparing the replication timing of ARS315 in its genomic location (Control, black) versus in its excised state as a chromatin circle (ARS315, blue) (**Fig. 3I**). Importantly, when comparing the ratio of ARS315 replication with a late replicating region on chromosome IV (Chr4, (35), see also **Fig. 1B),** the two replication profiles showed similar kinetics where ARS315 started replicating between 24 to 32min and DNA copy numbers increased. After ∼40min release, the region on Chr4 also started to replicate and DNA copy numbers decreased in both strains, consistent with earlier replication of ARS315 compared to Chr4. We note that the excised ARS315 chromatin circles did not reach the same maximum DNA copy number ratio at 40min as the Control genomic ARS315, suggesting that a fraction of chromatin circles did not replicate as efficiently as in its genomic context which could stem from the lack of stochastic firing of neighboring origins and passive replication when isolated from its genomic context. We also tested whether the process of recombination and purification affected chromatin accessibility at individual sites on the chromatin circles by restriction enzyme accessibility. Importantly, purified ARS305 and ARS313 chromatin rings (eluate) and isolated nuclei from the same strains before and after recombination displayed similar sensitivities to restriction endonucleases, suggesting essentially identical chromatin structures (**Fig. S3E-F**), as previously demonstrated for other chromatin domains purified by this approach^33^. We conclude that excision of the targeted origin from its genomic context does not interfere with the major chromatin properties of the origins and therefore also preserves the intrinsic replication state of the origin.

### MATAC-Seq and Psoralen-crosslinking Electron Microscopy show comparable single-molecule nucleosomal profiles on the multicopy ribosomal ARS domain

We next used a combination of M.CviPI/M.SssI (GpC/CpG-5-methyl-Cytosine-specific) and EcoGII (methyl-6-Adenine-specific) DNA methyltransferases, which catalyze preferential methylation of accessible DNA bases. Direct detection of methylated nucleotides by nanopore sequencing allowed us to generate single-molecule readouts of the chromatin accessibility states of individual ARS domains. In addition, we spiked an *in vitro* reconstituted nucleosomal array with 12x601 nucleosome positioning sequences(47) and a naked plasmid DNA control into the reaction to determine overall DNA methylation efficiencies and variability between individual biological replicates (**Fig. 4A**). To establish the assay and optimize the methylation conditions, we first focused on the multi-copy ribosomal ARS locus, which could be purified in large quantities and high yields as previously shown (**Fig. 4B** and (44)). The nucleosomal array revealed a methylation pattern that was consistent with highly positioned nucleosomes formed over the 601 sequences and intervening accessible linker DNA (**Fig. 4C-D**). In contrast, the methylation pattern of the purified rARS domain, as an example of a native chromatin domain, was much more irregular without equally spaced peaks (**Fig. 4E-F**). At the center of the individual nucleosomal rings, a highly methylated and thus accessible region was clearly visible that coincided with the annotated position of the rARS NFR. In addition, the flanking regions around the rARS displayed higher resistance against methylation, indicating nucleosomal protection. Importantly, a regular pattern of unmethylated DNA interspersed with highly accessible linker DNA was not as pronounced as for the nucleosomal array, suggesting a much higher level of heterogeneity at this locus (**Fig. 4E-F**).

**Figure 4.**
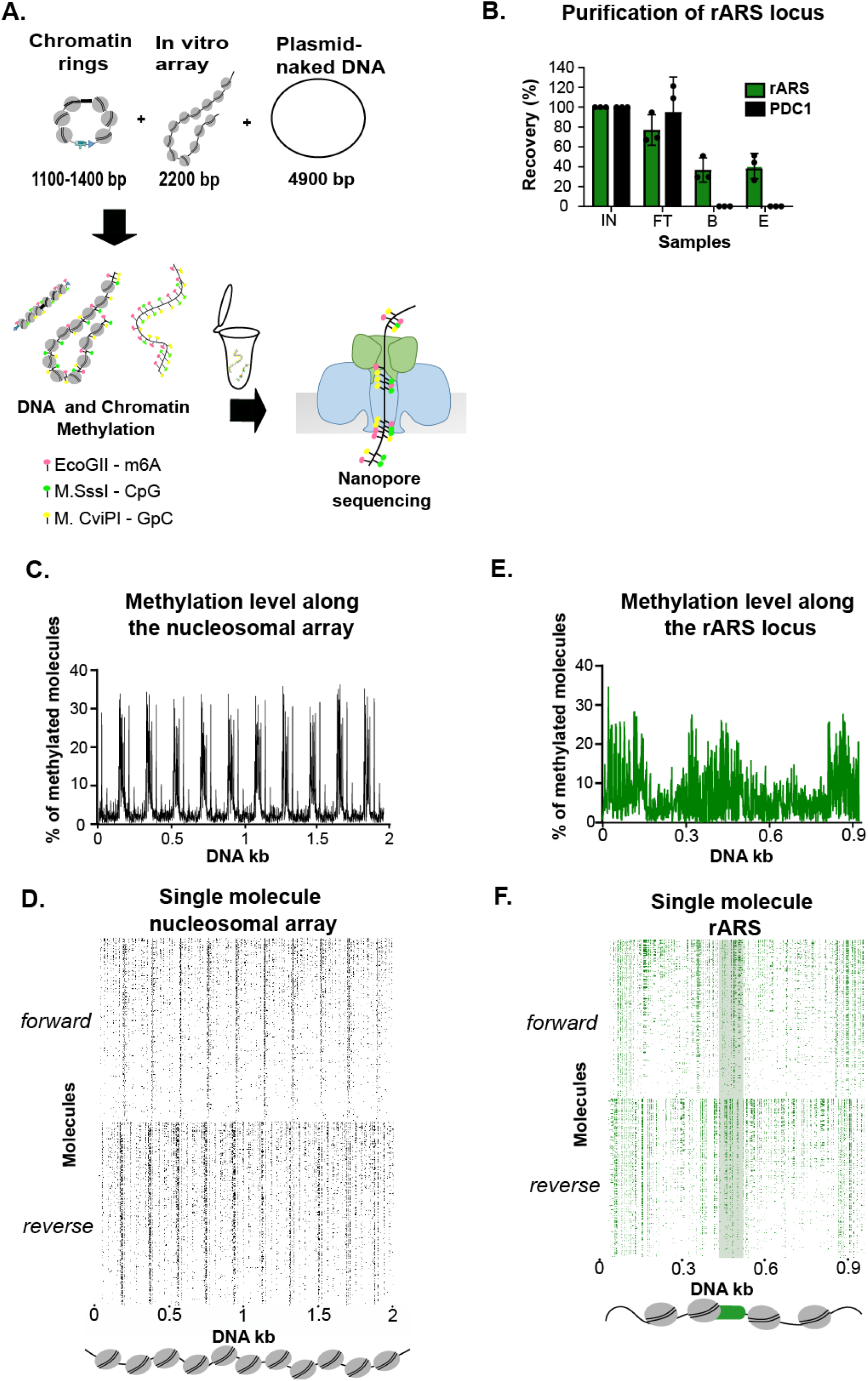
**A.** Experimental outline of single molecule **M**ethylation **A**ccessibility **TA**rgeted **C**hromatin loci sequencing assay (MATAC-seq). The purified chromatin rings and two spike-in controls; an *in vitro* assembled nucleosomal array and a naked plasmid were treated with m6A, CpG and GpC methyltransferases, which preferentially methylate only accessible DNA bases. DNA was isolated and the circular molecules were linearized with BsrGI and subjected to nanopore sequencing. **B.** LexA affinity purification was performed for strain Y84, where the multi-copy rARS locus is targeted by RS. DNA samples were taken (0.1% Input (IN), Flowthrough (FT), Beads (B), Eluate (E) from n=3 biological replicates and analyzed by qPCR. The enrichment of an unrelated region (PDC1) was analyzed side-by-side with the regions of interest. **C.** Average methylation profile of *in vitro* assembled nucleosomal array. MATAC-seq reveals the repetitive pattern of protected nucleosomal regions separated by narrow hyper-methylated linker sites. **D.** MATAC-Seq reads of the *in vitro* assembled nucleosomal array. Each row represents a single molecule displaying methylation events at near bp resolution. The reads have been ranked according to their methylation level (higher on the top and lower on the bottom) on both DNA strands. **E.** Average methylation profile on the native rARS locus reveals high accessibility at the centrally positioned ARS region. **F.** MATAC-Seq reads of the multi-copy rARS chromatin locus. Each row represents a single molecule displaying methylation events at near bp resolution. The reads have been ranked according to their methylation level (higher on the top and lower on the bottom) on both DNA strands. Identical number of reads were used per DNA strand and per sample (1560 reads) in MATAC-seq plots.

We benchmarked this single-molecule methylation nanopore dataset with psoralen-crosslinking electron microscopy (EM), considered as a gold standard approach to analyze single-molecule nucleosome configurations(43–45, 48). To this end, we incubated purified rARS chromatin rings with trimethylpsoralen and exposed to UV-A light, which is known to result in interstrand-crosslinked DNA in linker but not nucleosomal DNA. Thus, positions previously occupied by nucleosomes are visible in denaturing electron microscopy as single-stranded DNA bubbles, separated by double-stranded linker DNA that resist denaturation due to psoralen crosslinking (**Fig. S4A**). DNA was linearized such that the nucleosome-free ARS region was in the center of the molecule and the positions of the observed surrounding DNA bubbles were mapped in all molecules (**Fig. S4B**). Overall, we detected 143 molecules that strictly conferred to the expected size range of ∼1kb (**Fig. S4C**). Importantly, the majority of detected bubbles (∼70%) on the ribosomal ARS molecules displayed the expected size range of mononucleosomal bubbles (**Fig. S4D,** light blue area). Of note, the residual 30% of the bubbles showed smaller (< 135bp), intermediate (165 – 300bp) or di-nucleosomal sizes, presumably because of incomplete crosslinking of naked DNA due to sequence preferences for psoralen-crosslinking, protection by chromatin components other than nucleosomes or inefficient linker DNA crosslinking between two nucleosomes (**Fig. S4D**). Interestingly, the molecules revealed substantial heterogeneity in the number, position, and size of DNA bubbles, consistent with our methylation footprinting analysis (**Fig. S4E**). These data suggest the co-existence of a variety of chromatin states at this locus. Unlike MATAC-Seq, the EM dataset did not allow to orient individual molecules. Thus, we could only directly compare the accessibilities between the two methods at the two ends and the centrally positioned ARS of the linearized molecules. Importantly, overlapping the bulk profiles generated from psoralen-EM with the methylation footprinting data revealed highly concordant profiles between the high accessibility regions (high methylation and low frequency of nucleosomal bubbles), whereas the intervening regions showed higher nucleosomal protection (**Fig.S4F**). We conclude that MATAC-Seq is robust and provides high-resolution chromatin accessibility maps of specific chromatin domains.

### MATAC-Seq reveals major changes in origin chromatin accessibility between wildtype and CRE mutant strains

Having established a method to determine chromatin states with single-molecule resolution, we next wanted to obtain insights into the potential single-molecule differences of origin chromatin accessibility between the wildtype and CRE mutant strains. To this end, we used site-specific recombination to excise native chromatin rings encompassing a ∼1kb genomic region centered around the single-copy ARS305, ARS313, ARS315 and ARS316 replication origins in the wildtype, isw2Δ and ies6Δ mutant strains. We first validated that the targeted purification of the 4 single-copy replication origins via LexA-TAP showed comparable purification recovery efficiencies and enrichments between the individual strains and biological replicates (**Fig. S5A-D**), although we note that ARS316 chromatin domains in the ies6Δ mutant were less efficiently recovered in the final eluate than for the wildtype and isw2Δ purifications (**Fig. S5B**). Importantly, genomic mapping of MATAC-Seq reads on chromosome III confirmed a strong enrichment of reads overlapping with the targeted replication origins and low numbers of contaminating genomic reads (**Fig. S5E**).

For the single molecule analysis, we used Megalodon(49–51) to extract the CpG and Adenine-methylated bases from raw nanopore reads. We took into account an equal number of reads from forward and reverse strands and obtained a very high read depth for each origin and condition (984-1560 reads), with the exception of the ARS316 ies6Δ mutant, for which we recovered fewer reads (548 and 166) per replicate **(Fig. S5F)**. To reduce the noise from false positive detection of methylation, we compared the methylation density scores of an unmethylated to fully methylated naked plasmid and set specific thresholds for CpG and m6A methylation (**Fig. S6A-B**). Importantly, we also validated the significance of our datasets by comparing the wildtype average methylation patterns generated by MATAC-Seq across all reads with bulk accessibility profiles of recently published ChIP-Exo datasets of canonical histone H3, ORC4, ORC5 and MCM5 at these loci(52) (**Fig. 5A-D**). Notably, the H3-depleted ARS region of each origin was hyper-accessible and overlapped strongly with the ORC and MCM ChIP-Exo peaks (**Fig, 5A-D**) and the nucleosome-free region (NFR) of the Tvs1 gene promoter downstream of ARS315 showed pronounced accessibility (**Fig. 5B**), further confirming that MATAC-Seq can reliably identify open regulatory chromatin regions.

**Figure 5.**
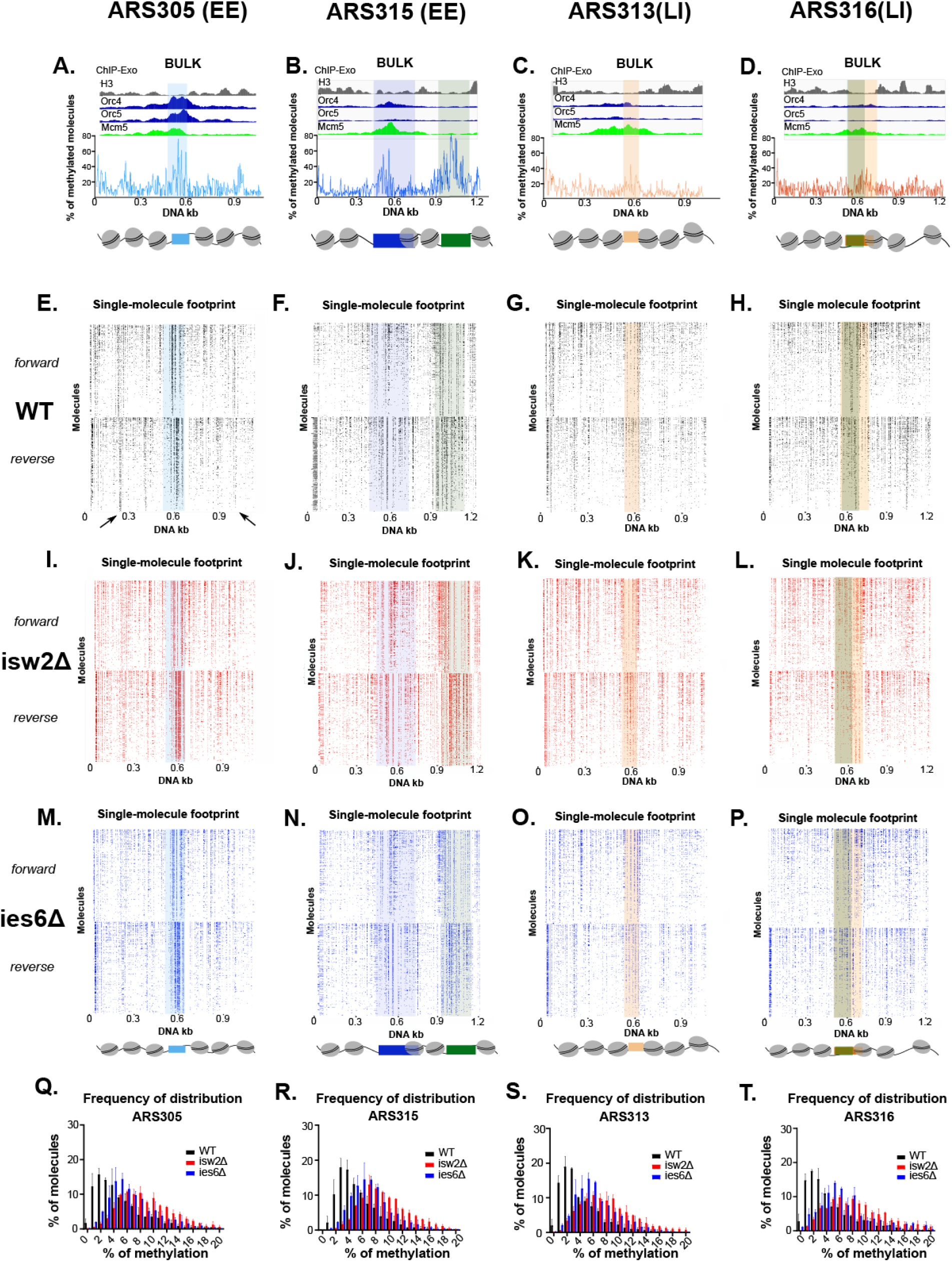
**A-D.** Average methylation profile around the replication origins (ARS305, ARS315, ARS313, ARS316) derived from MATAC-Seq recapitulates the known nucleosomal pattern derived from bulk ChIP-Exo analysis^55.^ **E-H.** Chromatin accessibility maps around the native ARS loci of wildtype strains revealing less nucleosomal ARS regions and high level of heterogeneity. The methylated DNA bases of each molecule are depicted as dots and the reads have been organized according to their methylation level (higher on the top and lower on the bottom) in both DNA strands. Identical number of reads were used for forward and reverse strands in MATAC-seq plots. **I-L.** MATAC-Seq chromatin accessibility maps of the ARS regions in the isw2Δ strains. Identical number of reads were used for forward and reverse strands in MATAC-seq plots. **M-P.** MATAC-Seq chromatin accessibility maps of the ARS regions in the ies6Δ strains. Identical number of reads were used for forward and reverse strands in MATAC-seq plots. **O-T.** Frequency of methylation distribution. Comparative analysis of frequency of methylation distribution in all replication origins between WT and CRE mutants shows an overall increase of unprotected regions in CRE mutants. Average and standard deviations are from n= 2 biological replicates.

We first ranked all single-molecule forward and reverse reads according to their total methylation level in the wildtype, isw2Δ and ies6Δ mutant (**Fig. 5E-P**). In the wildtype maps, multiple locations (for example at the –3 and +3 nucleosomal regions of ARS305) displayed strongly positioned nucleosomal footprints with nearly all individual reads exhibiting an accessible linker region between the nucleosomes (**Fig. 5E,** black arrows). However, most visible substructures exhibited considerable heterogeneity with a gradient in methylation levels along both strands, suggesting an overall diverse protein occupancy landscape likely generated by the co-existence of multiple chromatin states on individual molecules. Notably, the WT strain clearly showed a more unstructured and diffuse methylation pattern in the two LI ARS313 and ARS316 origins in comparison to the EE origins ARS305 and ARS315 (**Fig. 5E-H**). This is in line with the fact that the EE single-molecule maps showed higher accessibility in the NFR regions of the ARS (see shaded boxes at the center of the bulk MATAC-Seq profiles in **Fig. 5A-B** versus **Fig. 5C-D**), which likely facilitates a more regular phasing of nucleosomes and other surrounding chromatin factors.

Interestingly, the overall frequency distribution of the percentage of methylated bases was shifted towards higher methylation levels in the two mutants compared to the wildtype. This effect was consistent across all four origins and more pronounced in the isw2Δ mutant (**Fig. 5I-T**), suggesting that deletion of ISW2 has a stronger impact on nucleosomal occupancy/chromatin accessibility than deletion of IES6. To exclude the possibility that these differences derive from technical variations – e.g. different efficiencies of the methylation reaction - we took advantage of the spike-in naked plasmid and nucleosomal array to assess the methylation levels and allow normalization between individual experiments (**Fig. S6C-D**). Importantly, the spike-in naked plasmid also contained the ARS305 sequence, allowing us to directly compare the methylation efficiency of the same DNA sequence in the context of naked DNA versus native chromatin. The chromatinized molecules showed clear protection from methylation except for the central ARS position that displayed similar accessibilities between the naked plasmid and native chromatin context (**Fig. S6E**).

### An optimal NFR size between 100 and 115bp with intermediate accessibility allows for efficient origin replication

We next asked whether the increased accessibility in the CRE mutants is due to a uniform gain of accessibility across the complete ∼1kb region or whether specific genomic regions were affected more than others. To this end, we divided the origins into smaller bins containing regions that are predicted to align with the center of individual nucleosomes (ns) based on the histone H3 ChIP-Exo profiles (**Fig. 5A-D**). Additionally, we assigned intervening linker DNA (L) between nucleosomes as well as the ARS of all four origins (blue-shaded boxes) and the promoter region (prom) in ARS315 (green-shaded box) as regions of interest (**Fig. 6A-D**). Importantly, the methylation profiles of wildtype and CRE mutant strains were highly reproducible between the biological replicates (**Fig. S6F-I**), allowing us to pool the replicates and increasing the statistical power. After normalization of the methylation levels of wildtype and mutant strains, we determined whether and which genomic regions showed a significant change in methylation level globally across the whole origin (**Fig. 6A-D**) and locally across the assigned genomic features (**Fig. 6E-H and Fig. S7A-D**).

**Figure 6.**
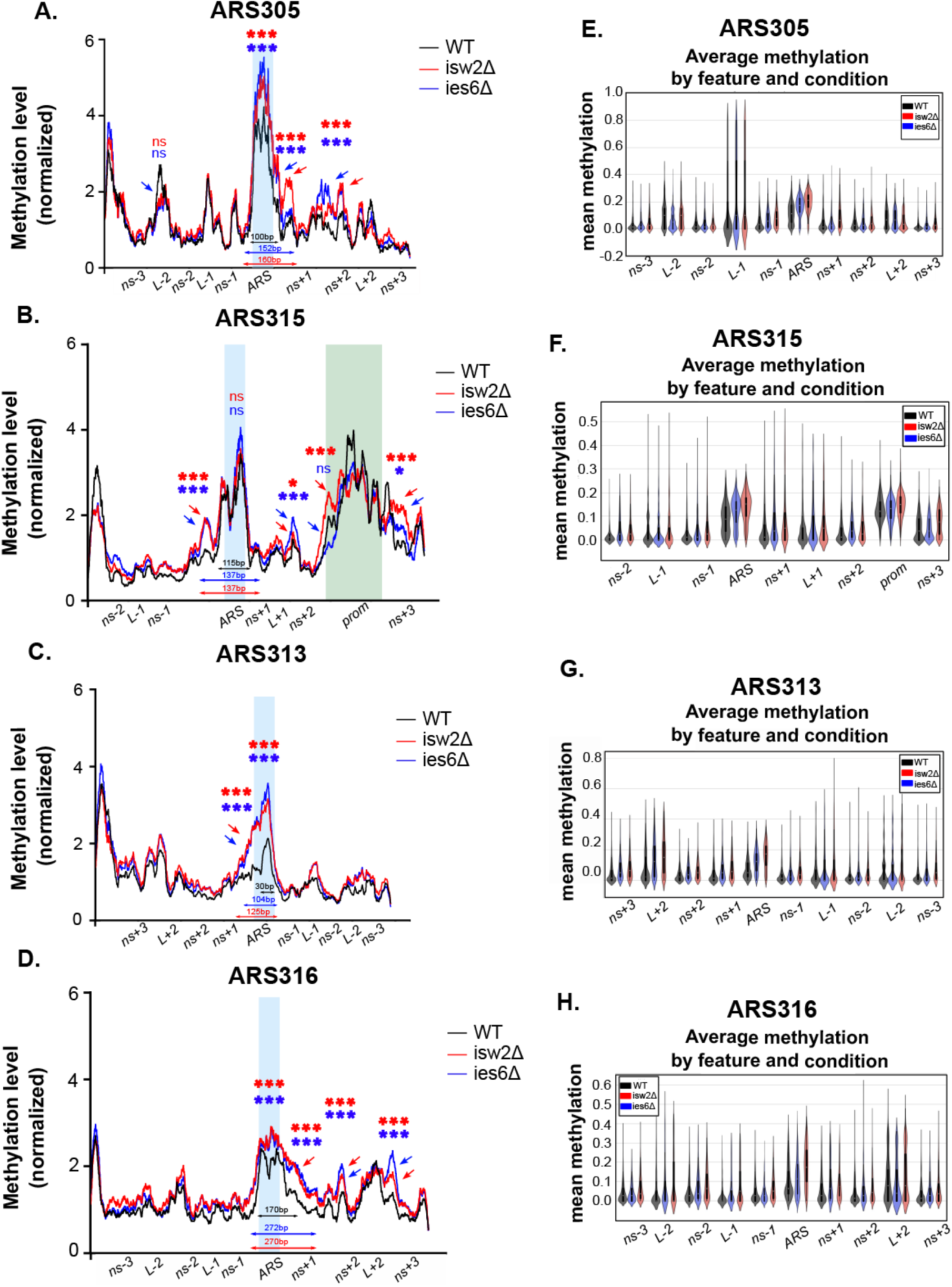
**A-D**. Comparative analysis of chromatin accessibility on replication origins between WT and CRE mutant strains shows statistically significant difference on specific features. The methylation level of each condition has been normalized to the maximum methylation value and smoothed using 30 bp window. The size of each the NFR of the ARS in wildtype and CRE mutants is indicated at the bottom using specific thresholds of mean methylation for each ARS (ARS305, ARS315 >1.2, ARS313 >1.4 ARS316 > 1.13). Statistical analysis performed between genomic bins representing nucleosomal positions (80bp long), linker regions (10-40bp) and the nucleosome-free ARS (100bp) and Prom in ARS315 (175bp). **E-H.** Comparative analysis of the average mean methylation of each specific features along the molecules of different replication origins between WT and CRE mutants. The size of ARS and the nucleosomal regions is 80 bp. The length of linker regions varies from 10 to 40 bp and the promoter region on ARS315 domain is 175 bp long. Average and standard deviations are from n= 2 biological replicates. (*, *** indicates statistical significance P < 0.05, or P < 0.001, respectively, by two-sided test).

One of the most striking changes observed consistently was that the accessibility of the ARS was significantly increased in both mutants. Importantly, this pattern was consistent for all replication origins (**Fig. 6E-H, ARS**), suggesting that both chromatin remodelers have overlapping functions in regulating ARS accessibility, either by directly sliding nucleosomes into the ARS or indirectly by establishing a more regular chromatin structure in direct proximity of the ARS. For each origin, we defined the size of the NFR based on a specific threshold of the normalized DNA methylation level (see also legend to **Fig.6**). Deletion of ISW2 increased the size of the NFR at the EE origin ARS305 from 100bp to 160bp and shifted the position of the linker between the +1 and +2 nucleosome further downstream (**Fig. 6A red arrows**), consistent with repositioning of the +1 nucleosome further away from the ARS. A similar broadening of the NFR to 152bp was observed for the ies6Δ mutant together with an increased accessibility between the +1 and +2 nucleosomes (**Fig. 6A, blue arrows**). Interestingly, the ARS315 (EE) origin showed similar changes for both mutants, with an increase in the size of the NFR on the side of the -1 nucleosome from 115bp in the wildtype to 137bp in the mutants (**Fig. 6B**). In addition, the isw2Δ mutant showed a broadening of the Tvs1 promoter NFR (**Fig, 6B, red arrows around prom**). The nucleosome upstream to the Tvs1 promoter was shifted towards the ARS, thereby reducing the spacing of the +1 and +2 nucleosomes located between the promoter and ARS (**Fig. 6B**). Similarly, the ies6Δ mutant showed an additional increase in accessibility at the L+1 linker region between the +1 and +2 nucleosome, suggesting that the positioning of these two nucleosomes was particularly affected. (**Fig. 6B, blue arrow at L+1**).

ISW2 and IES6 deletion also resulted in a strong increase in accessibility at the otherwise well-protected ARS of ARS313 (LI) and ARS316 (LI), but also led to an increased size of the NFR on the + sides of the two ARS (**Fig. 6C-D, red and blue arrows**). Interestingly, the ARS313 NFR significantly broadened from 30bp in the wildtype to 104bp in the ies6Δ mutant and to 125bp in the isw2Δ mutant (**Fig. 6C**). At the ARS316 origin, both isw2Δ and ies6Δ mutants broadened the NFR from 170bp in the wildtype to 270bp and 272bp in the CRE mutants, respectively. The enlargement of the NFR coincided with an increased accessibility downstream at two additional positions, consistent with a shift or sliding of the +1 and +2 nucleosomes in both mutants (**Fig. 6D**).

Together, these differential accessibility patterns of the CRE mutants align with changes of the EE and LI states of the origins. First, the EE origins both decrease replication efficiency upon ISW2 or INO80 deletion (**Fig. 1C-D**). In both cases, we observed increased accessibility as well as broadening of the ARS NFR to larger sizes than the wildtype length of 100bp (ARS305) and 115bp (ARS315), suggesting that the positioning or distance of the +1 and -1 nucleosomes might have a strong impact on the EE replication property of these origins. Second, the LI origin ARS313 advanced replication in the ies6Δ but not the isw2Δ strain (**Fig. 1E**). Intriguingly, the ies6Δ strain broadened the NFR to the same size range (104bp) as the wildtype NFR of the two EE origins ARS305 and ARS315 (100bp – 115bp), whereas the ARS313 NFR in the isw2Δ mutant slightly exceeded this range with a size of 125bp. Finally, no advancement of replication of the LI origin ARS316 was observed in both CRE mutants (**Fig. 1F**). Although overall accessibility and size of the NFR was also increased for both CRE mutants, the wildtype ARS316 NFR already showed a larger size of 170bp compared to the two EE origins, supporting the notion that specific chromatin features such as an optimal size of the NFR but not high accessibility of a locus per se can serve as a positive regulator of replication efficiency.

### Hierarchical clustering reveals a distinct chromatin state with an open ARS and an open +2 linker region that correlates with efficient origin activity

In order to leverage the power of our single-molecule datasets and identify groups of molecules with common chromatin accessibility states, we applied hierarchical clustering analysis on the molecules derived from the four replication origins in wildtype, ies6Δ and isw2Δ mutants. To this end, we used the same bins as in **Fig. 6** to divide the origins into distinct regions that are predicted to be nucleosome-free (ARS and Prom), nucleosomal regions (NS) and linker regions between nucleosomes (L) and calculated the average methylation level per feature and molecule. Based on these features, hierarchical clustering analysis of each origin produced a tree that was pruned to a height of 5 distinct clusters (**Fig. 7A-D**), so that each cluster presented a unique chromatin accessibility state. Importantly, splitting the reads that contribute to each of the 5 clusters according to biological replicates showed highly similar contributions from the two independent experiments, but distinct contributions from the experimental condition per cluster (wildtype, ies6Δ and isw2Δ mutants) (**Fig. S7E**). Additionally, each cluster constituted a similar number of reads derived from forward and reverse strand (**Fig. S7F**), indicating that the clustering algorithm did not segregate by DNA sequence and thus potentially different numbers of methylation sites present on the two DNA strands. Based on the resulting heatmaps for each origin (**Fig. 7A-D**), we assigned for each cluster whether a given feature showed high, intermediate and low methylation levels to illustrate the relative chromatin accessibility state along the origins (**Fig. 7A-D, cartoons on the right**). We then determined in a bar graph presentation the relative percentage of reads that contributed to each cluster from wildtype, ies6Δ and isw2Δ mutant conditions (**Fig. 7E-H**).

**Figure 7.**
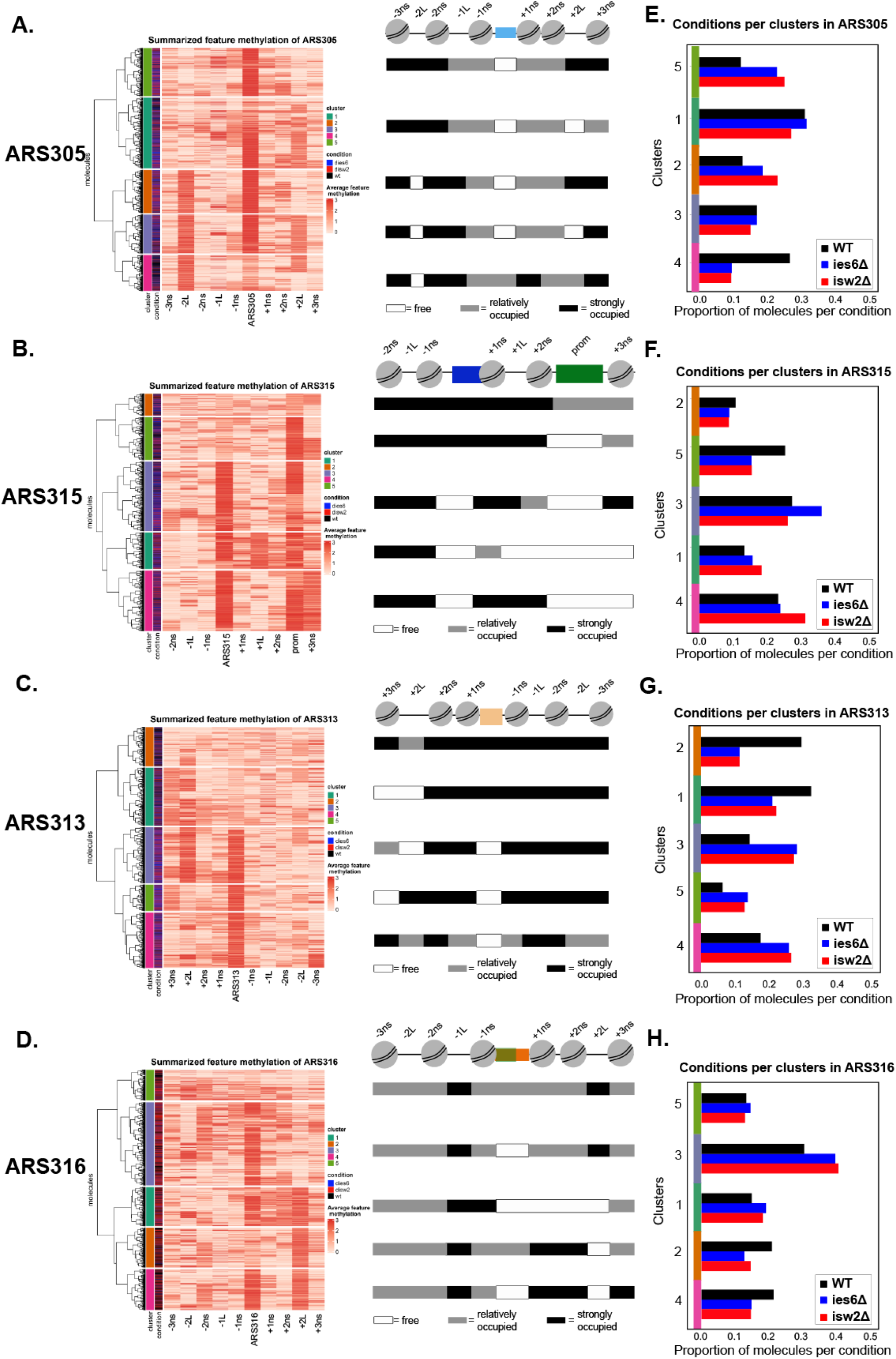
**A-D.** Heatmaps showing hierarchical cluster analysis of the summarized methylation levels per feature along the single ARS molecules. For each origin, the dendrogram was cut after 5 clusters. The cartoons on the right represent a visual representation of the associated chromatin accessibility state. Regions with high methylation level are considered as NFR and depicted as white. Regions with medium methylation level are considered as relatively occupied nucleosomal sites and depicted as grey and the low methylated features are considered as fully nucleosomal and represented as black. **E-H.** The plots show the proportion of the reads per condition and cluster. The total number of reads has been normalized to one.

MATAC-Seq revealed a disperse set of dynamics and cluster occupancies between wildtype and CRE mutants at the investigated EE and LI origins. In ARS305 (**Fig. 7E**), cluster 1 is the main cluster in both wildtype and CRE mutant strains and shows an open ARS site and open +2L surrounded by relatively well positioned nucleosomes. Interestingly, cluster 4, characterized by an open -2L, but less accessible ARS, is the second largest cluster in wildtype cells. This chromatin state is less frequently observed in both ies6Δ and isw2Δ mutants, whereas more reads contribute to clusters 2 and 5 in the CRE mutants. The common feature of clusters 2 and 5 is the protection of the +2L, suggesting that nucleosomes have shifted and occupy this large linker region of the origin. Given that the CRE mutants increase the accessibility and reposition nucleosomes, this may indicate that the +1NS, +2 NS and +3NS are not as precisely positioned in these two clusters and therefore breach more frequently into this linker region.

In ARS315 (**Fig. 7F**), cluster 3 represents the most abundant chromatin state in the wildtype cells containing both an open ARS and open Prom-site flanked by well positioned nucleosomes. Interestingly, cluster 5, characterized by a closed ARS but open Prom-site is the second largest cluster in wildtype cells, which challenges previous models that an open ARS region is a necessary and uniform requirement of an EE origin based on bulk experiments(17, 21). This state is less frequently observed in the CRE mutants, where more reads contribute to cluster 3, 4 and 1 instead of cluster 5. As these clusters display a generally higher accessibility of the surrounding chromatin of ARS and Prom-sites, these data suggest that the strong positioning of the +1NS, +2NS and +3NS are instructive to efficient replication, as observed in the other EE origin ARS305.

ARS313 is an LI origin that advanced replication most significantly in the ies6Δ CRE mutant (**Fig. 1E**). In wildtype cells, clusters 1 and 2 containing a closed ARS as a common feature contribute most reads to the population (**Fig. 7G**), suggesting that the strong ARS occupancy is a limiting feature at this origin. Intriguingly, the CRE mutant strains contributed less to the clusters 1 and 2 but shifted a significant portion of reads to clusters 3, 4 and 5 that all displayed an open ARS. Most strikingly, the largest cluster in the ies6Δ mutant was cluster 3, characterized by an open ARS and a large highly accessible +2L region, which represents a highly similar chromatin state as cluster 1 in the EE origin ARS305, the most frequent cluster in wildtype cells (**Fig. 7E**). This result would also fit with the data from the second EE origin ARS315 if one considers the nucleosome-free Prom-site as a corresponding large linker region between the +2 and +3 nucleosomes (**Fig. 7F**).

Finally, the second LI origin ARS316 did not display changes in replication timing between WT and CRE mutant cells (**Fig. 1F**). The largest number of MATAC-Seq reads in wildtype cells corresponded to cluster 3 containing an open ARS flanked by relatively well but not strongly occupied regions (**Fig. 7H**). This cluster was even further enriched in the CRE mutant samples at the cost of clusters 2 and 4. As the common feature of clusters 2 and 4 was again a large open +2L region and the mutants showed this state less frequently, this result provided further support that the combination of an open ARS with an optimal size of 100-115bp together with an open +2L region flanked by well positioned nucleosomes is a chromatin state that supports or at least highly correlates with early and efficient replication of these four origins. Together, our MATAC-Seq data allowed us to extract key chromatin features of stochastic origin activation previously masked in common population-based techniques.

## Discussion

To date, the chromatin landscape of EE and LI origins have mostly been investigated by bulk assays showing that ARS are located in a nucleosome-free region surrounded by well-positioned nucleosomes. Here, we developed MATAC-Seq as a single-molecule method combining the enrichment of targeted loci with a chromatin accessibility profiling assay, thereby providing locus-specific chromatin occupancy maps at unprecedented resolution and coverage. This provides a solid basis for detailed characterization of alternative and rare chromatin states at a given locus of interest, addressing current limitations of genome-wide single-molecule techniques to study chromatin accessibility(42, 43, 53, 54).

Evaluation and comparison of MATAC-Seq with complementary methods is an essential step to validate our results. Averaging the single-copy locus methylation profiles could recapitulate bulk ChIP-Exo profiles and clearly identifies NFRs of replication origins as well as neighboring gene promoters (**Fig. 5A-D and Fig. 6**). Additionally, we evaluated the MATAC-Seq profiles with an alternative single-molecule method by psoralen-crosslinking EM analysis on the multi-copy rARS locus (**Fig. S4F**). Since this approach requires specialized equipment as well as high input amount of DNA that we only obtained from a multi-copy gene locus, this strengthens the usefulness and feasibility of MATAC-Seq to monitor targeted single-copy domains. However, the method is currently limited to a handful of selected loci, for which genetic manipulations to insert RS/LEXA sites are required and therefore restrict the genome-wide throughput of the method. Extending MATAC-Seq by targeting and enriching larger chromosomal regions of several 10 - 100 kilobases combined with the advantage of third generation long-read sequencing technologies is a future direction to be explored.

MATAC-Seq provides the first investigation of EE and LI origins after their *ex vivo* isolation from native chromosomes. Interestingly, our results show substantial level of heterogeneity at all origins investigated. There are molecules at both EE origins consistent with a fully nucleosomal configuration, but also subpopulations completely accessible without nucleosomes as well as intermediate states with well-defined surrounding chromatin structure (**Fig. 5 and Fig. 7**). How such high levels of heterogeneity functionally impact the stochastic activation of replication origins is an important question. To this end, we applied MATAC-Seq to CRE mutant strains of ISW2 and INO80, implicated in nucleosome sliding and eviction(55–57). Interestingly, both isw2Δ and ies6Δ strains showed decreased replication efficiency at the EE origins (**Fig. 1C-D**) which was accompanied with higher chromatin accessibility at the ARS (**Fig. 6A-B**), suggesting that these chromatin remodelers are at least partially responsible to establish a well-positioned + or -1 nucleosome, thereby restricting the size of the NFR to an optimal size of 100-115bp to become an efficient substrate for replication initiation at EE origins. In contrast, the accessibility of the LI origin ARS316 was also strongly increased in both mutants (**Fig. 6D**) without apparent consequences to replication efficiency (**Fig. 1F**). This suggests that a general increase of accessibility/loss of chromatin structure is not sufficient for efficient origin activation, but rather the presence of key neighboring chromatin components at specific locations determines replication efficiency. We found that the size of the NFR may represent one such critical parameter as in wildtype cells, the two LI origins showed either a very small NFR of 30bp at ARS313 or a large NFR of 170bp at ARS316 (**Fig. 6C-D**) compared to an intermediate size of 100-115bp at the EE origins ARS305 and ARS315, respectively (**Fig. 6A-B**). At all four origins, the CRE mutant strains increased their relative NFR size compared to the wildtype condition, but strikingly, only the ies6Δ strain at the LI origin ARS313 increased its size to 104bp and therefore fell in the same range as the wildtype NFR sizes of 100-115bp of the two EE origins (**Fig. 6C**). Accordingly, the ies6Δ mutant increased origin firing efficiency by 20% (**Fig. 1E**), suggesting that an optimal size of the NFR between 100-115bp may represent a critical determinant of efficient origin replication.

Hierarchical clustering analysis revealed large heterogeneities between alternating states of accessible and non-accessible regions for each molecule and origin (**Fig. 7**). For consistency, we classified and divided the molecules for each origin into 5 distinct clusters that represent different chromatin states. This choice was instructed by the fact that the resulting 5 clusters did not show major biases in their contribution from biological replicates and/or the ratio of forward/reverse strands (**Fig. S7E-F**) and thus likely represent biological and not technical differences in chromatin accessibility. However, there are clear examples of individual clusters that show two populations of distinct methylation states at specific regions (e.g the +2L region in cluster 4 of ARS305 or the +3NS in cluster 5 of ARS315). Thus, we cannot exclude the presence of additional biologically meaningful subclusters in the data and further refinement of this analysis may provide more insights into the variability of origin chromatin. Nevertheless, the choice of 5 clusters per origin revealed clear differences in the relative contribution of reads from wildtype and CRE mutant cells (**Fig. 7E-H**). Comparing the four origin datasets, an interesting finding is that the combination of an open ARS with a long and open +2 linker region flanked by well positioned nucleosomes appears as a chromatin state that supports or at least highly correlates with early and efficient replication. The mechanistic link of how such an asymmetric chromatin state could favor the bidirectional and thus symmetric process of replication initiation is an open question, but one plausible explanation could be that the open +2 linker region provides an additional binding site in the right distance and orientation to the ARS for the binding of additional factors, for example the replication initiation factors Cdc45, Sld2 or Sld3 known to be limiting for early and efficient origin firing(58). To address this question, an interesting perspective would be to perform proteomic analysis of the purified chromatin domains in the CRE mutant strains, as we have recently established in the wildtype condition (manuscript under revision).

Clearly, the small number of replication origins in this study limits the possibility to draw global conclusions but gives us the opportunity to strongly correlate and further dissect the functional relationship between the chromatin accessibility landscape and the replication profiles of these four ARS loci. In future work, we expect MATAC-Seq as a broadly applicable long-read single-molecule method to study the functional importance of heterogeneity of chromatin states, which is a major driver for cellular plasticity implicated in development as well as many human diseases such as cancer (59–64).

## Supporting information

Table of Plasmids

Table of Primers

Table of Yeast strains

## Acknowledgments

This work was funded by the German Research Foundation (Deutsche Forschungsgemeinschaft, DFG) via the Collaborative Research Cluster SFB1064. J.K., A.S and M.L. were supported by the SNF Project Grant 310030_189206. We thank Bihter Özdemir Aygenli and Till Bartke for providing us with the nucleosomal array. We thank Adam Burton and Maria-Elena Torres-Padilla for critical reading and helpful comments on the manuscript.

## Author Contributions

Conceptualization and methodology, A.C., T.S., J.K., A.S., E.K., A.S. M.L. and S.H.; data curation, formal analysis, and software, A.C., T.S., K.H., and S.H.; visualization, A.C., K.H. and S.H.; funding acquisition, project administration, supervision, and resources, S.H. investigation and validation, A.C., M.W., T.S., K.H., J.K., A.S., H.U., E.K., M.T., M.W., M.L., M.L., A.S., and S.H. writing (original draft), S.H.; writing (review and editing), A.C., M.W., T.S., K.H., J.K., A.S., H.U., E.K., M.T., M.W., M.L., M.L., A.S., and S.H.

## Declaration of Interests

The authors declare no competing interests.

## Material & Methods

### Culture of yeast strains expressing the recombination cassette

Yeast strains competent for recombination were inoculated to OD600 0.2 in 2 L YPR medium at 30 °C and 200 rpm. At OD600 1.0, cells were arrested in G1-phase by α-factor (50ng/ml) and, simultaneously, recombination was induced by addition of galactose to a final concentration 2% (w/v) for 2h at 30 °C. Cells were harvested by centrifugation (6000 rpm, 10 min, 4 °C), the supernatant was discarded, and the cell pellets resuspended with water and collected in a sealed 25ml syringe. After pelleting the cells (3000 rpm, 12 min, 4 min) and discarding the supernatant, the cells were extruded into liquid nitrogen, broken in small ‘spaghetti’ pieces and stored at -80 °C for later use.

### Specific chromatin domain isolation with affinity purification

A commercial coffee grinder (Gastroback, 42601) was pre-cooled by grinding 30 g of dry ice for 30sec. The resulting powder of dry ice was discarded. Appropriate amount of frozen cell pellets (3-4 g of multi copy rARS circles or 5-6 g of single copy ARS circles) were mixed with ∼ 60 g of dry ice and ground in the coffee mill. The coffee grinder is run for 10x30 sec, with short intervals in between. Occasional tapping with a spatula against the outside of the mill prevents ground cells from sticking in a layer to the inside wall of the grinder. The fine powder of ground yeast was transferred into a plastic beaker until the dry ice evaporated. For chromatin methylation and nanopore sequencing experiments, frozen pellets of each yeast strain were ground separately and then pooled together for subsequent affinity purification. After evaporation of dry ice, the powder is dissolved in 0.75 ml of cold buffer MB (20 mM Tris-HCl pH 8.0, 200 mM KCl, 5 mM MgAc, 0.5 % Triton X-100 (w/v), 0.1 % Tween-20 (w/v)) with 1x Protease and Phosphatase Inhibitors (Protease and Phosphatase Inhibitor Cocktail 100x Thermo Fisher Scientific) per 1 g of ground yeast cells. The cell lysate was transferred into 2ml low-binding reaction tubes (Eppendorf) and was cleared from cell debris by centrifugation for 30 min in a microcentrifuge at 16.000 x g and 4 °C. Supernatant containing the diluted chromatin rings was transferred into 2ml low-binding tubes and incubated with IgG-coupled magnetic beads (500ul slurry of beads per 4 g of frozen cell pellet)(44). DNA and protein samples (0.1% and 0.05%, respectively) were obtained from the supernatant (Input IN) before the addition of magnetic beads. Supernatant was incubated with the beads for 2 h in a rotating wheel at 4 °C. Beads were separated from the supernatant using a magnetic rack and samples for DNA and protein analysis were collected from the resulting Flow Through (FT) (0.1% and 0.05%, respectively). The beads were washed 5 times with 1ml cold buffer MB with 1x Protease inhibitors and one final wash with 1ml cold buffer MB without protease inhibitors. For each washing step the beads were incubated on a rotating wheel for 10min. LexA-Chromatin ring complexes were released by proteolytic cleavage of the LexA-TAP fusion protein by overnight incubation with 2ul (purification of a single yeast strain) or 15ul (purification of all yeast strains together) 6xHis-tagged recombinant TEV protease in a total volume of 300 μl (purification of a single chromatin domain) or 400 μl (purification of pooled chromatin domains) in MB without protease inhibitors. Finally, beads were separated from the eluate containing the chromatin rings. DNA and protein samples were taken from beads (B) and final elution (E) (0.1% and 0.05%, respectively).

### Specific chromatin domain isolation with affinity purification – *DNA analysis*

DNA samples from the purification process were supplemented with H_2_O to a final volume of 100 μl. Additionally, 1,13 ng of plasmid DNA (K71) were added in each sample as control for the DNA extraction efficiency. 100 μl of IRN buffer (50 mM Tris–HCl pH 8, 20 mM EDTA, 500 mM NaCl) and 2μl RNAse A (10mg/ml) were added to the DNA samples. After incubation at 37 °C for 1 h, 5 μl Proteinase K (10mg/ml) and 10 μl of SDS 20% were added and incubated for an additional 1 h at 56 °C. Subsequently, 200µl Phenol:Chloroform:Isoamyl Alcohol (25:24:1, v/v) was added, followed by 2 x 10 sec thorough vortexing. The organic and aqueous phases were separated by centrifugation for 7 min at 21.000 x g. The supernatant was transferred to a fresh 1.5 ml tube containing ethanol (2.5 : 1) and 2 µl glycogen (10mg/ml). The tube was left at -20 °C overnight. The solution was centrifuged with 21.000 x g at 4 °C for 45 min. The supernatant was discarded and 150 µl of 70 % ethanol was added to the pellet. After another centrifugation step with 16.000 x g at 4°C for 10 min, the resulting DNA pellet was dried at room temperature for 10 min. DNA samples were digested with BsrGI restriction endonuclease (NEB) in a total volume of 35 μl and DNA content analyzed by qPCR.

### Specific chromatin domain isolation with affinity purification – *Protein analysis*

Protein samples were resuspended in 50 µl 1x SDS sample buffer and boiled at 95 °C for 3 min. Whole cell extracts were separated by electrophoresis, transferred onto polyvinylidene difluoride membranes (Immobilon®-P PVDF Membrane, Sigma-Aldrich) and blocked in 5 % skimmed milk dissolved in 0.05 % Tween/PBS (PBST) for 1 h at room temperature. Membranes were incubated with primary antibody Peroxidase Anti-Peroxidase Soluble Complex antibody produced in rabbit for detection of TAP-tagged proteins (Sigma-Aldrich, P1291-500UL) 1:1000 dilution in 5 % skimmed milk/PBST overnight at 4 °C followed by washing in 0.1%Tween/TBS. Membranes were developed by chemiluminescence.

### Chromatin and DNA methylation

The protocol of enzymatic methylation treatment of chromatin was adopted and optimized from (42). The final elution from pooled chromatin purifications (∼200 ng total DNA in 400 μl) was supplemented with 100ng of *in vitro* assembled nucleosomal array and 1 μg of naked plasmid DNA (K112), which were used as control of methylation efficiency and nucleosome detection. Chromatin and naked DNA were treated first with 200 U of EcoGII and 200 U of M.CviPi (NEB), SAM (S-adenosylmethionine) (NEB) at 0.6 mM and sucrose at 300 mM, and then incubated at 30 °C for 7.5 min. After this incubation, additional 100 U of each enzyme and 0.05 mM SAM were added and the incubation was continued for 7.5 min at 30 °C. Subsequently, 120 U of M.SssI (NEB), 10 mM MgCl_2_ and 0.05 mM SAM were added to the reaction with continued incubation at 30 °C for 7.5 min. Additional 0.05 mM SAM were added and further incubation at 30 °C for 7.5 min was carried out. Methylation reaction was stopped by Stop Buffer to 1:1 volume (20 mM Tris-HCl pH 8.5, 600 mM NaCl, 1% SDS, 10 mM EDTA), followed by DNA extraction as described below. All reactions were performed in low-binding reaction tubes (Eppendorf).

### Chromatin and DNA methylation - *DNA extraction*

The methylated fragments were incubated with 2 μl RNAse A (10mg/ml) for 1 h at 37 °C and 5 μl Proteinase K (10mg/ml) followed by 1 h incubation at 56 °C. Subsequently, 200 µl of Phenol:Chloroform:Isoamyl Alcohol (25:24:1, v/v) was added, followed by 2 x 10 sec thorough vortexing. The organic and aqueous phases were separated by centrifugation for 7 min at 21.000 x g. The supernatant was transferred to a fresh 1.5 ml tube containing ethanol (2.5 : 1) and 2 µl glycogen (10mg/ml). The tube was left at -20 °C overnight. The solution was centrifuged with 16.000 x g at 4 °C for 45min. The supernatant was discarded and 150µl of 70% ethanol was added to the pellet. After another centrifugation step with 16.000 x g at 4°C for 10min, the supernatant was again discarded. The DNA pellet was dried at room temperature for 10min. The dried pellet is then resuspended in 40µl H_2_O and digested with BsrGI (NEB) in a total volume of 50 μl for 2 h at 37 °C. After digestion, DNA was cleaned using PCR purification kit (ThermoFisher Scientific). In the final step, DNA was eluted into nuclease-free H_2_O in a total volume of 50 μl and used for library preparation.

### Library preparation

For MATAC-seq, the recovered DNA was converted into libraries using Ligation Sequencing Kit 1D (Oxford Nanopore Technologies, SQK-LSK109) following the manufacturer’s instructions. Nanopore sequencing was carried out on R9.4 MinION flowcells (Oxford Nanopore Technologies) for up to 24 h.

### Nanopore basecalling

Megalodon (including megalodon 2.4.0, guppy 5.0.11, minimap 2.17 and res_dna_r941_min_modbases-all-context_v001.cfg) was used for base calling and detection of CpG and m6A methylation. Thresholds for binarizing the methylation states were set as follows: we generated density plots of methylation scores of 10 million methylation sites analyzed on unmethylated and fully methylated plasmid samples, respectively, and determined a score cutoff - separately for m6a and CpG methylation - that yields in particular high specificity **(Fig. S5 A-B)**

### Analysis of nanopore sequencing data

Binary count data was normalized to average methylation levels of the spiked-in nucleosome array controls, respectively. We then pooled replicates from the same condition, calculated mean methylation levels per position **(Fig. 4A-D)** and used the paired Wilcoxon-rank-sum test to extract p-values. Heatmaps in Fig. 4 show raw, binary data. To summarize methylation levels per feature (**Fig 4 E-H**), mean normalised methylation was calculated for each molecule and feature. For clustering, we scaled normalized average feature methylation to zero mean and a standard deviation of one, and calculated the distance matrix as 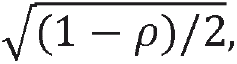 where ρ is the Spearman’s correlation coefficient between molecules. We applied hierarchical clustering (using R’s hclust and the ward.D2 method), and cut the dendrogram after 5 clusters. Heatmaps in Fig X show average feature methylation per molecule, as well as cluster assignment and condition. For the histograms in Fig Y, we normalized total number of reads per condition to 1 and display the proportion of reads per condition and cluster. Scripts and binary input data are available under https://github.com/KarolineHoller/nanopore_methylation. (Repository will be made accessible for reviewers upon request.)

### Replication timing

Yeast cells were inoculated at OD600 0.2 in 2 L YPD medium at 30 °C and 200 rpm. At OD600 0.6, cells were arrested in G1-phase by α-factor (50ng/ml) for 2 h. Samples of 1 ml culture for flow cytometry analysis were obtained every 30 min (0, 30, 60, 90, 120 min). After 2h G1-arrest, cells were released into S-phase by addition of 125 U of Pronase (Sigma-Aldrich, 53702-25KU) and potassium phosphate buffer to final concentration of 20 mM. Samples for flow cytometry and quantitative PCR (1 ml and 4.5 ml, respectively) were collected every 8 min for 48 min.

### Replication timing – *quantitative PCR*

The samples were supplemented with 500 μl of 1% sodium azide solution (w/v) in 0.2 M EDTA, washed once with water (3000 x g, 3 min, 4 °C) and the resulting yeast pellets were snapfrozen in liquid nitrogen. For DNA extraction, the pellets were resuspended into RINB (50 mM Tris HCl pH 8, 0.1 M EDTA, 0.1% (v/v) beta mercaptoethanol) with Zymolyase to a final concentration of 2% (w/v) and were incubated for 1 h at 37 °C and 700 rpm shaking. After addition of 2μl RNAse A (10mg/ml), spheroblasts were incubated for 1 h at 37 °C, followed by the addition of 5 μl Proteinase K (10mg/ml) and 10 μl SDS (20 %) at 55 °C for 1 h. DNA was isolated by phenol-chloroform extraction (1:1) followed by overnight ethanol precipitation as described above. DNA pellets were suspended in 50 μl of H_2_O and 10 μg of DNA was digested by EcoRI restriction endonuclease (NEB) at 37 °C for 1 h. DNA samples were diluted 1:10 with H_2_O and analyzed by qPCR.

### Replication timing – *Flow cytometry*

Cells were centrifuged at 16000 x g for 2 min. The supernatant was discarded and 1ml of cold 70 % ethanol was added slowly drop-by-drop with gentle agitation. The fixed cell suspensions were stored at 4 °C until further use. 500 – 600 μl of the samples were transferred into a clean 1.5 ml reaction tube, centrifuged and the resulting pellets resuspended with 300μl 50 mM Na-citrate and 0.1mg/ml RNAseA. After 2 h incubation at 50 °C, 3 μl Proteinase K (10mg/ml) was added (1:100) and another 2 h incubation at 50 °C was followed. 30 μl of each sample were mixed with 170 μl of 50 mM Na-citrate and 0.5 μM Sytox Green (S7020, ThermoFisher). Prior to the FACS analysis, the samples were briefly sonicated (Bioruptor, 5min with 30 sec ON and OFF intervalls) to detach cell clumps before proceeding with the analysis.

### Restriction enzyme accessibility assay of linear, recombined and affinity-purified chromatin

Cells were grown to OD600 0.8 in 50 ml of YPR medium at 30 °C and 200 rpm. 25 ml of yeast culture were centrifuged (3000 x g, 10 min, 4 °C), washed twice with H_2_O, snapfrozen in liquid nitrogen and were stored at -20 °C for later use. Cell pellets were washed three times with Buffer A (15 mM Tris-HCl pH 7.4, 0.2 mM Spermine, 0.5 mM Spermidine, 80mM KCI) and 1x Protease Inhibitors (16 rpm, 2 min, 4 °C) and then resuspended in 350 μl of Buffer A x Protease Inhibitors and ∼450 μl glass beads (1mm, BioSpec Products). The buffer solution should be enough to cover the beads by a thin layer. The cell pellets were vortexed thoroughly for 2x10 min with interval break of 5 min. To collect the cell lysates, the bottom and cap of microtubes were pierced with a hot needle and placed in a 15ml tube. After centrifugation (130 x g, 1 min at 4°C in a microcentrifuge), the glass beads remained in the microtubes. The crude cell lysates, collected in the 15ml tubes, were transferred into new 1.5ml microtubes. The 350 μl suspension was supplemented with 2 mM MgCl_2_ to the total volume of 400 μl. To achieve restriction digestion at chromatin loci by the indicated enzymes, the crude nuclei, or the purification elution containing the chromatin rings, was split into different reaction tubes for each restriction enzyme tested. Digestion was performed for 60 min at the optimal reaction temperature using different amounts of the respective restriction endonuclease (10 or 50 U and 50 or 100 U as indicated). The reaction was terminated by adding IRN buffer (1:1). Samples were treated with 2μl RNAse A (10mg/ml) followed by 1 h incubation at 37 °C and 5μl Proteinase K (10mg/ml) followed by 1 h incubation at 56 °C. DNA was isolated by phenol-chloroform extraction (1:1) followed by overnight ethanol precipitation. DNA was linearized by restriction enzyme digestion overnight at 37 °C in a final volume of 50μl. The DNA samples were subjected to indirect end-labeling Southern blot analysis.

### Southern Blot

Nucleic acids from genomic DNA were separated on a 1% agarose gel and blotted onto a positive Nylon membrane (Amersham Hybond™-N, GE) by capillary transfer in 1 M Ammonium acetate. DNA probes for hybridization were generated using the RadPrime DNA labeling system (Invitrogen) with incorporation of [α−32P]dATP (Hartmann Analytik) according to the instructions of the manufacturer. Images were acquired with the Typhoon FLA 7000 imaging system.

### Psoralen crosslinking

The 400ul eluate with the purified chromatin rings was transferred into two wells of a 24well culture plate. Each sample was supplemented by 10ul Trimethylpsoralen (TMP) (0.2 mg/ml in Ethanol), mixed and incubated in dark on ice for 5 min. Samples were positioned 5 cm away from a 366-nm ultraviolet bulbs in a Stratalinker and then irradiated for 5min. Additional incubation of 6min, 7min, 8min were performed and before each incubation additional 10μl TMP were added. After irradiation, the samples were transfered to microtubes, respective wells washed once with 200μl IRN and combined with the samples (total volume 400μl). After treatment with RNaseA (at a final concentration of 0.33 mg/ml for 2 h at 37 °C), Proteinase K and SDS were added to a final concentration of 0.33 mg/ml and 0.5% and incubation was continued for 4 h at 55 °C. DNA was extracted with phenol/chloroform and precipitated overnight at 4 °C. The DNA was then digested with 3ul NcoI restriction enzyme (NEB) in 23μl total reaction volume.

### Denaturing spreading and analysis by electron microscopy (EM)

The procedure was performed as recently described(65). For denaturing spreading, the reaction consisted of 2.0ul formamide, 0.4ul glyoxal and 2ul of chromatin rARS locus, was incubated for 10 min at 42°C in a thermo-mixer and subsequently transferred to an ice water bath. After this denaturation step, the reaction was spread onto carbon-coated 400-mesh magnetic nickel grids over a water surface using the benzyldimethylalkylammonium chloride (BAC) method. Following this spreading procedure, the DNA was platinum coated by platinum-carbon rotary shadowing (High Vacuum Evaporator MED 020; Bal-Tec) to render it electron dense. The grids were scanned using a transmission electron microscope (Tecnai G2 Spirit; FEI; LaB6 filament; high tension ≤ 120 kV) and pictures were acquired with a side mount charge-coupled device camera (2,600 × 4,000 pixels; Orius 1000; Gatan, Inc.). The images were processed with DigitalMicrograph Version 1.83.842 (Gatan, Inc.) and analyzed using ImageJ64.

## Figures

**Figure S1.**
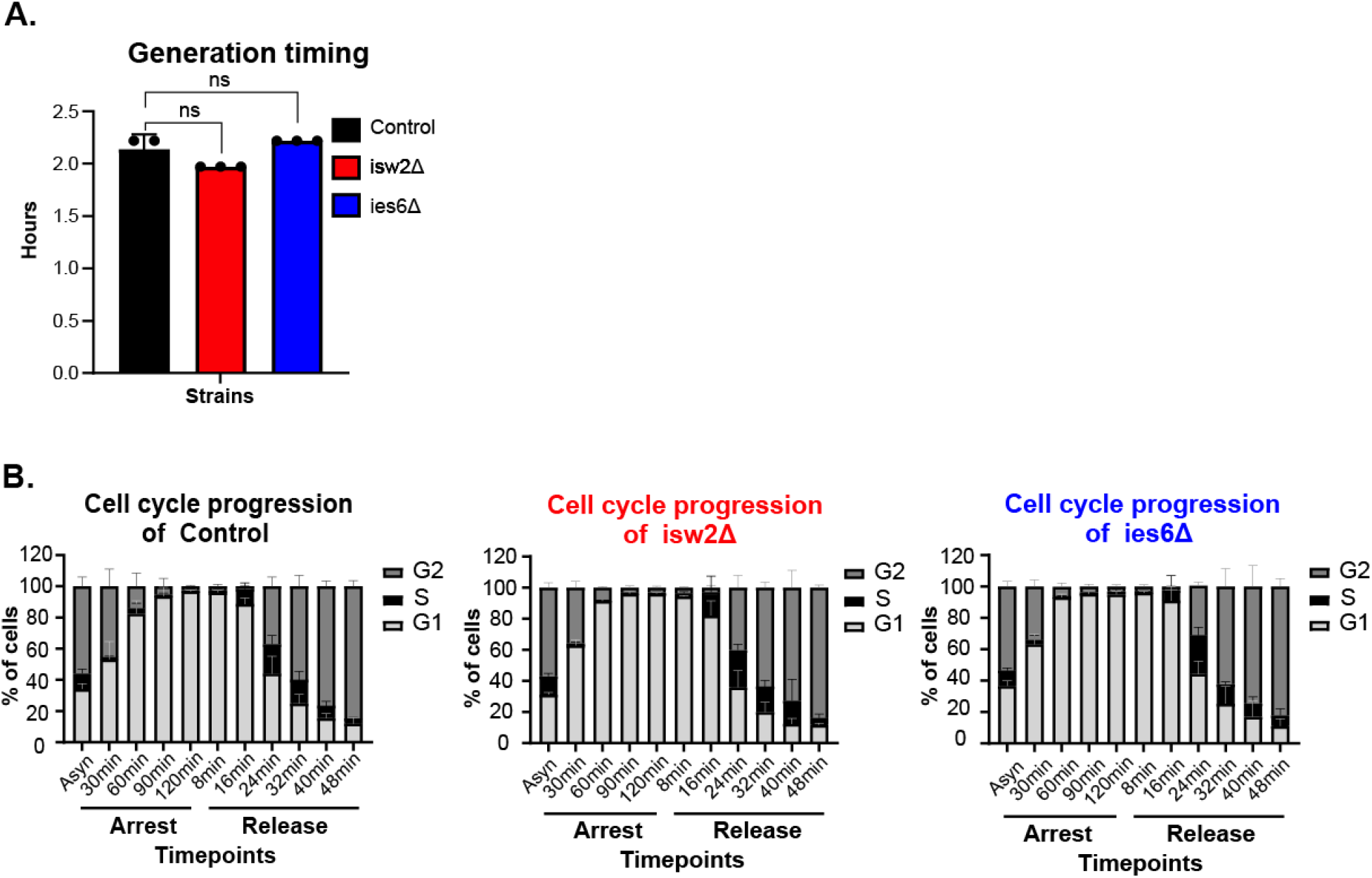
**A.** Generation timing of Y66 (control), Y104 (isw2Δ) and Y136 (dies6Δ). The strains were grown in YPD medium and OD600 measurements were taken every 360 sec by a multi-plate reader. **B.** Cell cycle progression of control and mutant strains. The cells were arrested by 50 mg/ml α-factor and released into replication by 125U pronase. The FACS samples were taken at the indicated timepoints and side-by-side with the DNA samples subjected to qPCR analysis in Fig.1C.

**Figure S2.**
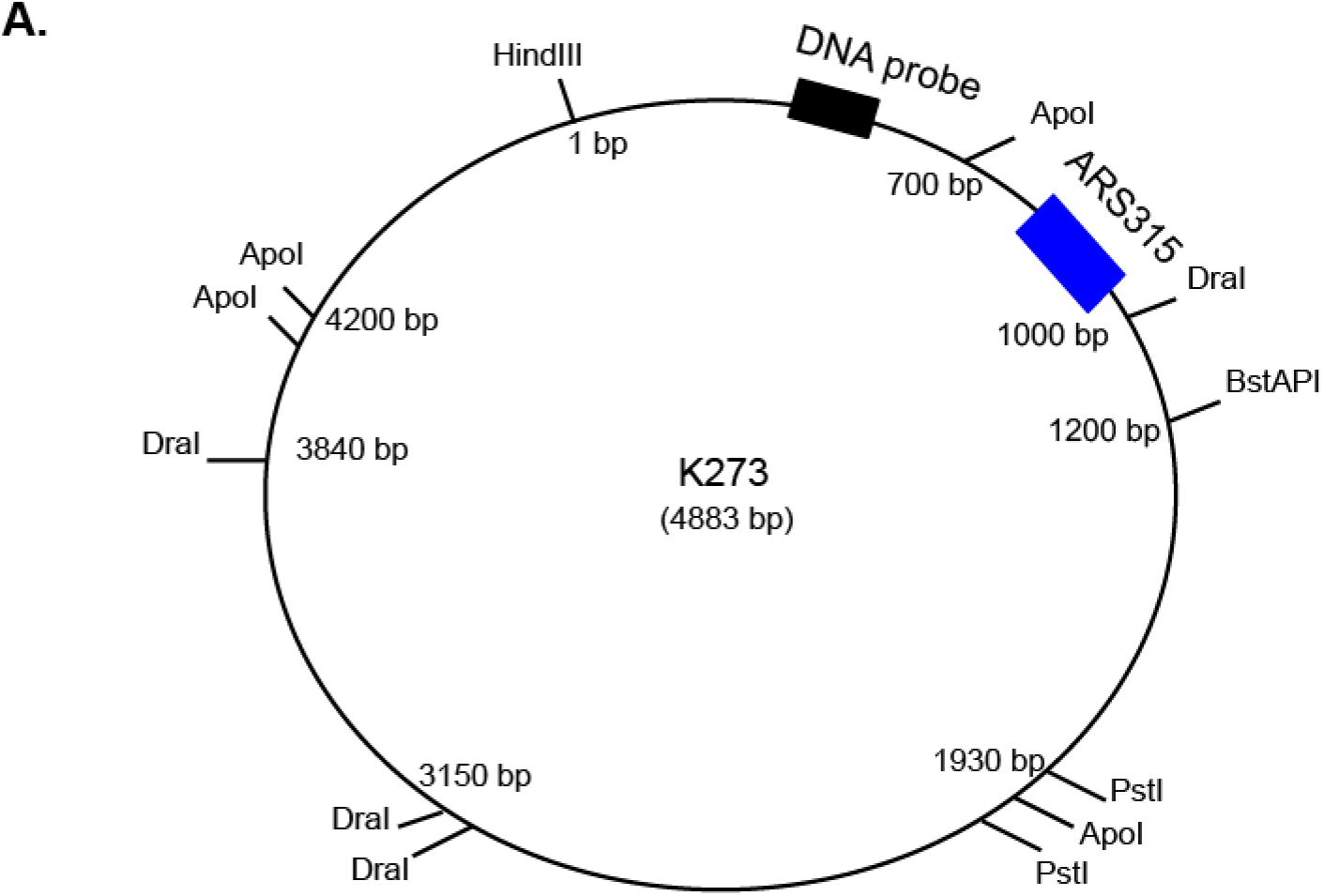
**A.** Schematic presentation of plasmid K273 containing the DNA sequence of ARS315 locus used as spike in control of the REA analysis of ies6Δ-ARS315 strains. The shared DNA sequence between the plasmid and the chromatin locus is highlighted in yellow. The black rectangle indicates the position of the DNA probe used for the Southern blot and the blue rectangle indicates the position of the ARS315 locus. On the map it is shown the distance between the restriction sites using the HindIII site as the starting point. Similar to the native chromatin locus, the plasmid was initially digested by BstAPI, DraI and ApoI, separately, and later all the samples were digested by HindIII and PstI (secondary digestion). The expected detectable DNA fragments according to the two digestion steps and the position of the probe are 1200 bp (for BstAPI and HindIII), 1000bp (for DraI and HindIII) and 700bp (for ApoI and HindIII). The larger fragments 4110 bp (in BstAPI) and 1300bp (in ApoI) indicate inefficient digestion of HindIII. All the fragments deriving from the plasmid are noted by asterisks in Fig. 2D.

**Figure S3.**
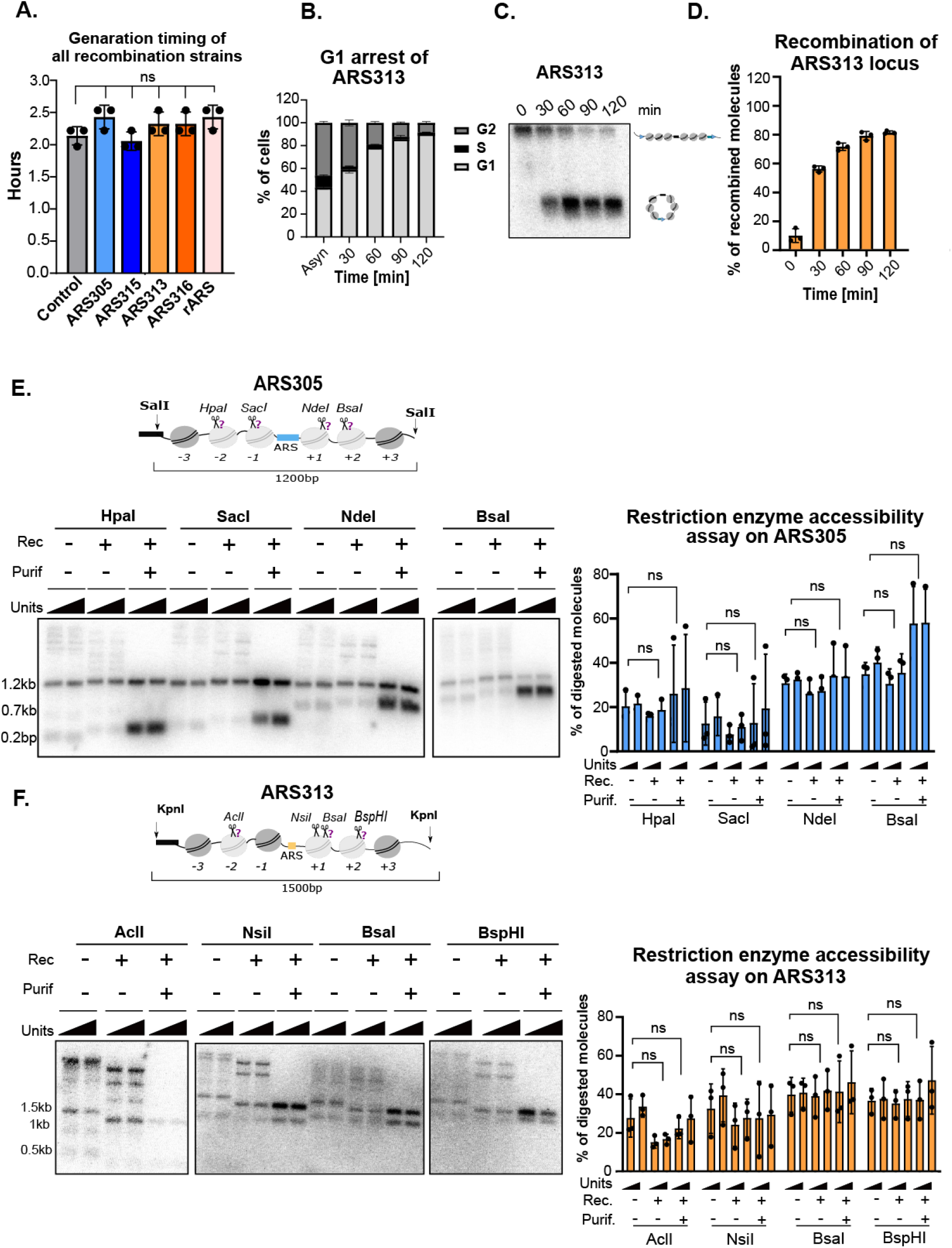
**A.** Generation timing of yeast strains competent for recombination (Y65-ARS305, Y91-ARS315, Y94-ARS313, Y69-ARS316, Y84-rARS) and control (Y66-without RS/LexA). The strains were grown in YPD medium and OD600 measurements were taken every 360 sec by a multi-plate reader. **B.** Cell arrest in G1-phase. The strain Y94-ARS313 was grown in YPR medium and arrested with 50 mg/ml α-factor. Samples for FACS analysis were taken at the indicated timepoints and stained by Sytox red to monitor the distribution of G1, S and G2-phase in both profiles. **C.** Recombination kinetics of ARS313 loci. The strain Y94-ARS313 and Y66-control (see Fig.3B) were grown in YPR medium to logarithmic phase and arrested in G1 phase by addition of α-factor (50ng/ml) and recombination was induced by the addition of 2% galactose. Samples were taken at indicated timepoints. DNA was isolated and linearized by BsrGI (in Y66 Fig.3B) and by NcoI (in Y94) and subjected to Southern blot analysis. The positions of unrecombined and recombined molecules are shown on the right. **D.** The histogram shows the results of Southern blot as percentage of recombined chromatin locus. **E-F.** Comparative REA analysis in chromosomal, recombined and purified ARS305 and ARS313 locus. Nuclei (before and after recombination) and chromatin rings (after purification) from yeast strains Y65 (ARS305), Y94 (ARS313) were isolated and digested with increasing amounts of the indicated restriction enzymes (scissors). DNA was isolated, digested with SalI in ARS305, KpnI in ARS313 and subjected to indirect end-labeling Southern blot analysis with the radioactively labeled probe. Top of each origin: schematic representation of each ARS locus with restriction sites used to probe chromatin structure (black rectangles) and restriction sites for isolation of DNA fragments for subsequent Southern blot analysis. The histogram shows the results of Southern blot quantification as a percentage of digested chromatin locus. Mean and standard deviations of all plots in Fig.S3 are from n= 3 biological replicates (ns, indicates no significant statistical difference P > 0.05, unpaired t-test).

**Figure S4.**
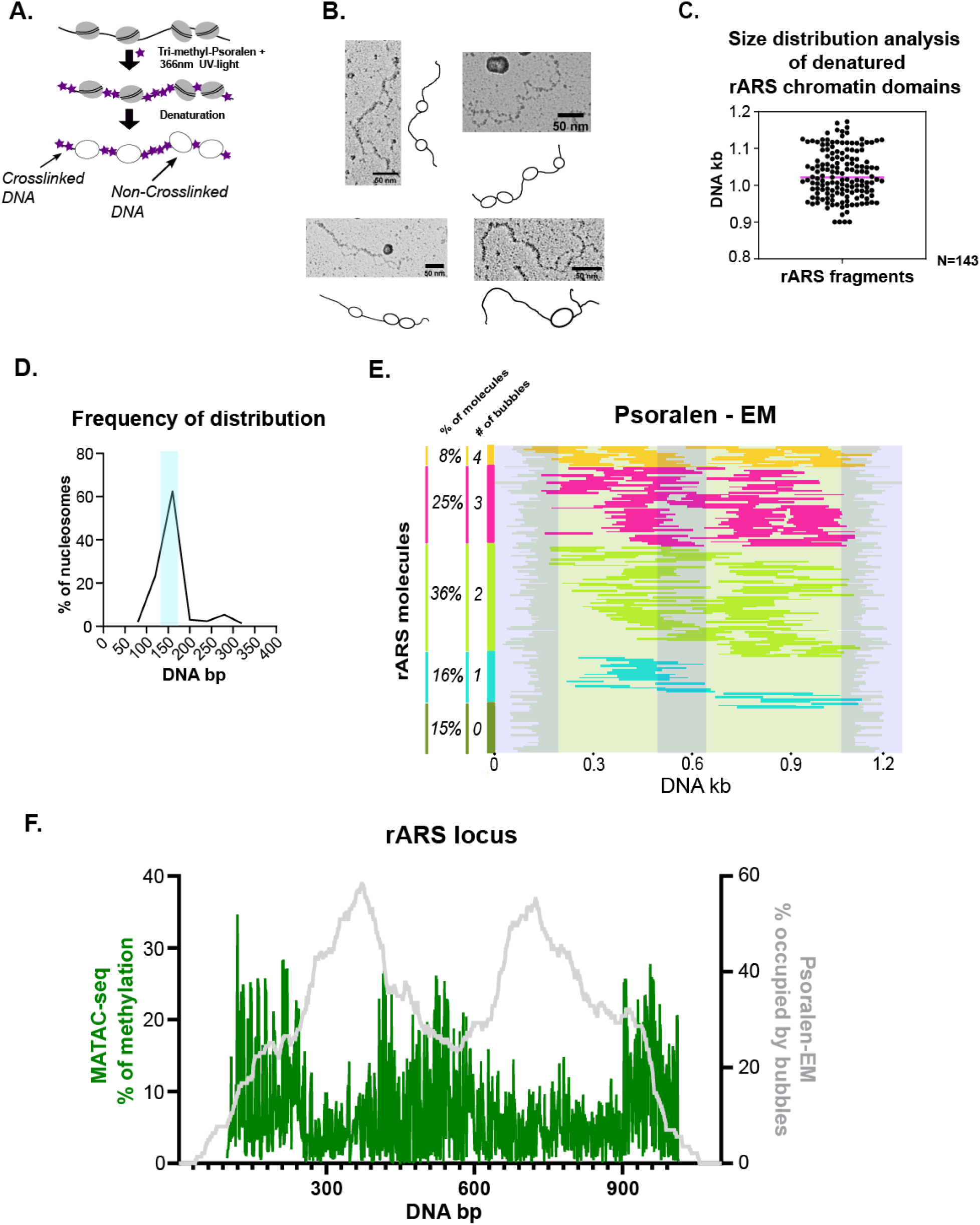
**A.** Schematic representation of the principal of the psoralen crosslinking assay. **B.** Determination of nucleosome positions at the rARS locus by single molecule EM analysis. Purified rARS chromatin rings were psoralen-crosslinked. After DNA isolation, the rings were digested by NcoI and then subjected to denaturing spreading followed by EM analysis. 143 molecules were analyzed by measuring the size and the number of each nucleosomal bubble. Representative electron micrographs for molecules containing different numbers of nucleosomes are shown on the left. **C.** Size distribution of the purified and denatured rARS fragments as analyzed by electron microscopy. The expected length of the rARS locus is 1046 bp +/-10% The pink line indicates the median. **D.** The plot shows the size distribution of the nucleosomal bubbles. The expected nucleosomal size is indicated in light blue. **E.** Five different groups of rARS molecules according to the number of nucleosomes (0, 1, 2, 3 or 4) and its coverage amongst the whole population. Each molecule has been aligned to its symmetric center position. The shadowed areas indicate the middle and the two ends of the rARS molecules. **F.** Averaging nucleosome profile of 143 unoriented rARS molecules. The plot shows the probability of a nucleosome in a specific locus capturing the less protected rARS origin and the two most prominent neighboring nucleosomal sites which is comparable to the averaging methylation profile derived from MATAC-seq.

**Figure S5.**
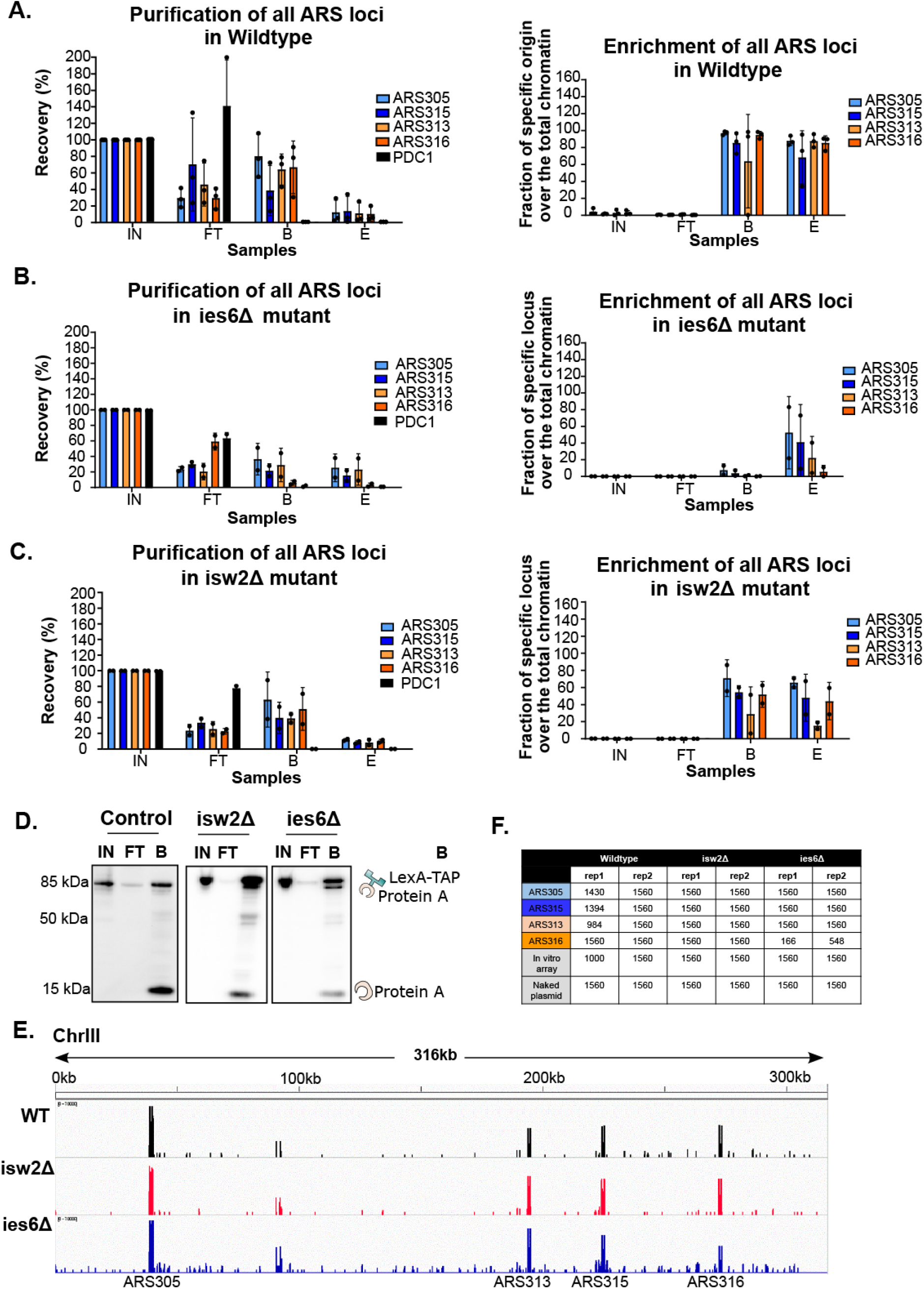
**A**. LexA affinity purification process was performed for all the WT strains together, which were then subjected to MATAC-Seq assay. DNA samples were taken (0.1% IN, FT, B, E) from n=2 analyzed by qPCR. The enrichment of an unrelated region (PDC1) was tracked side-by-side with the regions of our interest. The plot on the left shows how much of the total DNA was originated by the chromatin locus of our interest. Given that the size of yeast genome is 12 kb, and the length of a chromatin ring is ∼1 kb, the fold enrichment ratio of the specific origin to PDC1 was used to define the enrichment of an origin in the DNA samples. **B-C**. The same strategy was used to analyze the DNA samples of the CRE mutant strains (ies6Δ and isw2Δ). **D.** Protein analysis of LexA affinity purification process. The protein samples (0,5% IN, FT, B, E) were collected in parallel with the previous DNA samples and loaded to an SDS gel in order to monitor the LexA protein during the purification by Western blot. Samples were taken from WT, ies6Δ and isw2Δ. **E.** Enrichment of the targeted purified loci on ChrIII after MATAC-seq performance. **F.** Table indicating the number of reads per replicate and condition that have been used for MATAC-Seq analysis.

**Figure S6.**
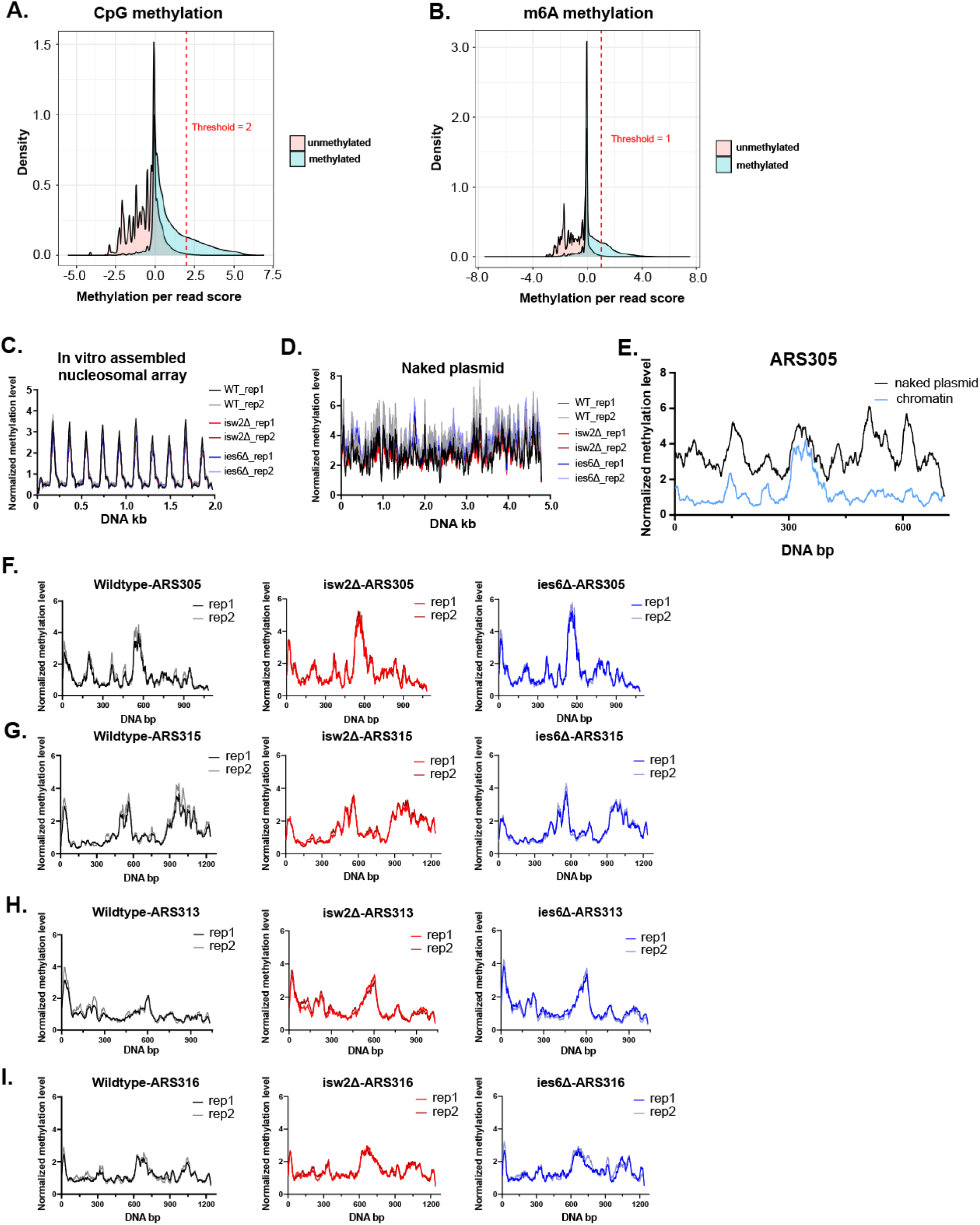
**A-B.** Methylation density comparison between a CpG/m6A methylated and an unmethylated plasmid. The dashed line indicates the threshold that was used to distinguish between signal and noise of methylation on the Nanopore sequencing data. **C-D** The plots show the methylation pattern on the in vitro nucleosomal array and naked plasmid (controls) of both biological replicates after normalization to maximum methylation. **E.** Comparative analysis of the same DNA sequence on ARS305 between naked plasmid and chromatin. The unprotected ARS domain of chromatin shows similar level of accessibility with the naked DNA. Their methylation level differs on the surrounding nucleosomal regions. **F-I.** Methylation pattern on native chromatin domains at ARS loci for both biological replicates. For all the plots in Figure S6, the methylation level has been normalized and smoothened using a 30bp window.

**Figure S7.**
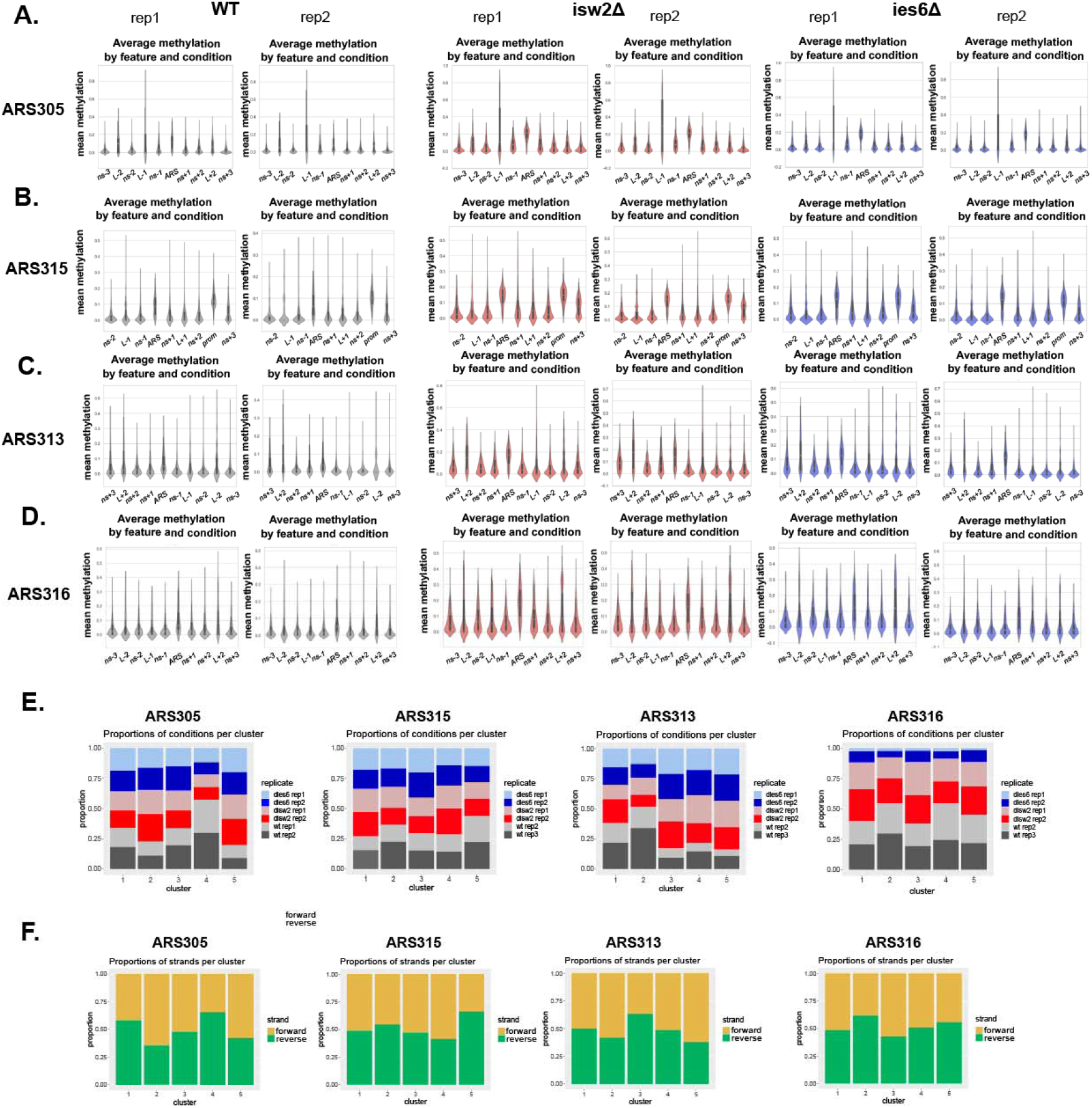
**A-D.** Average mean methylation of each feature around the replication origin chromatin domain of both biological replicates in WT, isw2Δ and ies6Δ strains. The size of each feature is similar to **Fig.6 E-H**. **E.** The plots show the read distribution of wildtype and CRE mutants per cluster. **F.** the plots show the proportion of forward and reverse DNA strands per cluster for each replication origin.

## Notes

### Competing Interest Statement

The authors have declared no competing interest.

