## Supplementary material for "Single molecule MATAC-seq reveals key determinants of DNA replication origin efficiency": Table of Plasmids

| | | |
|---|---|---|
| K004 (pM49.2) | (Griesenbeck et al., 2004) | Plasmid pM49.2 is a derivative of pABX22, and has been modified by addition of a LexA-binding cluster juxtaposed to an RS element. |
| K005 | Yeast expression vector for constitutive expression of LexA-TAP under control of TEF2 promoter and inducible expression of R Recombinase under control of GAL1-10 promoter; LEU2 selection marker framed with RS sites<br><br>(Hamperl et al., 2014) | |
| K009 | E. coli/yeast shuttle vector used for genomic integration of CYC1 LexATAP GAL1-10 RecR expression cassette by recombination in URA3 locus<br><br>(Hamperl et al., 2014) | |
| K18 | | pBlueSkript SK (+) |
| K94 | Plasmid containing wildtype ARS305 locus in pBlueScript backbone<br>(Matthias Weiß et al unpubiblished) | HindIII/PstI cut amplicon (primer 0016/0017) from yeast gDNA into K18 |
| K95 | Plasmid containing wildtype ARS316 locus in pBlueScript backbone | EcoRI/PstI cut amplicon (primer 0020/0021) from yeast gDNA into K18 |
| K102 | Vector with ARS305 +-2 nucleosome sequence next to lexA/RS sites<br><br>(Matthias Weiß et al unpubiblished) | Gibson assembly of two fragments (primer 0119/0120 with K94 as template + HpaI/XhoI cut backbone from K0004) |
| K103 | Vector with ARS305 +-3 nucleosome sequence next to lexA/RS sites<br><br>(Matthias Weiß et al unpubiblished) | Gibson assembly of two fragments (primer 0121/0122 with K94 as template + HpaI/XhoI cut backbone from K0004) |

| | | |
|---|---|---|
| K112 | Vector for yeast transformation in order to modifiy ARS305 locus and insert RS sites next to NS+-2<br>(Matthias Weiß et al unpubiblished) | Gibson assembly of two fragments (primer 0046/0047 with K94 as template + primer 0048/0049 with K102 as template) |
| K113 | Vector for yeast transformation in order to modifiy ARS305 locus and insert RS sites next to NS+-3<br>(Matthias Weiß et al unpubiblished) | Gibson assembly of two fragments (primer 0050/0051 with K94 as template + primer 0052/0053 with K103 as template) |
| K116 | Vector for yeast transformation in order to modifiy ARS316 locus and insert RS sites next to NS+-3<br>(Matthias Weiß et al unpubiblished) | Gibson assembly of two fragments (primer 0074/0075 with K95 as template + primer 0076/0077 with K106 as template) |
| K139 | E. coli/yeast shuttle vector used for genomic integration of pCYC1 LexA-TAP GAL1-10 RecR expression cassette by recombination in 500bp homology region from K121 of yeast chromosome I, LEU2 selection marker framed with RS sites (two mutations in the lexA gene that stop binding to lexA binding site: V11A, N171D)<br><br>(Matthias Weiß et al unpubiblished) | AscI/NheI cut amplicon (primer 0270/0271 template K121) inserted with AscI/NheI cut amplicon (primer 0272/0273 template K009). Resulting plasmid was cut with NheI/PacI and inserted with NheI/PacI cut amplicon (primer 0274/0275 template K009). |
| K173 | Plasmid containing wildtype ARS313 locus in pBlueSkript backbone | EcoRI/HindIII cut amplicon primers 319and 320 from gDNA into K18 |
| pM49.2_ARS313+-1 | Vector with ARS313+-1 nucleosome sequence next to lexA/RS sites | Gibson assembly of two fragments primers 321and 322 with K173 as template and HpaI/XhoI cut backbone from K004 |

| | | |
|---|---|---|
| pM49.2_ARS313+-2 | Vector with ARS313+-2 nucleosome sequence next to lexA/RS sites | Gibson assembly of two fragments primers 331and 332 with K173 as template and HpaI/XhoI cut backbone from K004 |
| pM49.2_ARS313+-3 | Vector with ARS313+-3 nucleosome sequence next to lexA/RS sites | Gibson assembly of two fragments primers 333and 334 with K173 as template and HpaI/XhoI cut backbone from K004 |
| K168 | Vector for yeast transformation in order to modify ARS313 locus and insert the RS sites next to NS+-1 | Gibson assembly of two fragments primers 346and 347 with pM49.2ARS315+-1 as template and primers 348 and 349 with K173 as template |
| K167 | E. coli/yeast shuttle vector used for genomic integration of pCYC1 LexA-TAP GAL1-10 RecR expression cassette by recombination in 500bp homology region from K121 of yeast chromosome I, LEU2 selection marker framed with RS sites<br><br>(Matthias Weiß et al unpubiblished) | Insert from K009 (NsiI/BlpI) cloned into K139 |
| K169 | Vector for yeast transformation in order to modify ARS313 locus and insert the RS sites next to NS+-2 | Gibson assembly of two fragments primers 350and 351 with pM49.2ARS315+-2 as template and primers 352 and 353 with K173 as template |
| K170 | Vector for yeast transformation in order to modify ARS313 locus and insert the RS sites next to NS+-3 | Gibson assembly of two fragments primers 366and 367 with pM49.2ARS315+-3 as template and primers 368 and 369 with K173 as template |
| K174 | Plasmid containing wildtype ARS315 locus in pBlueSkript backbone | EcoRI/HindIII cut amplicon primers 323and 324 from gDNA into K18 |
| pM49.2_ARS315+-2 | Vector with ARS315+-2 nucleosome sequence next to lexA/RS sites | Gibson assembly of two fragments primers 775and 776 with K174 as template and primers 771 and 772 with K004 as template |
| pM49.2_ARS315+-3 | Vector with ARS315+-3 nucleosome sequence next to lexA/RS sites | Gibson assembly of two fragments primers 777and 778 with K174 as template and |

| | | |
|---|---|---|
| | | primers 771 and 772 with K004 as template |
| K238 | E.coli/yeast shuttle vector used for genomic integration of pTEF2 LexA-TAP GAL1-10 RecR expression cassette by recombination in 500bp homology region from K121 of yeast chromosome I, LEU2 selection marker framed with RS sites<br><br>(Matthias Weiß et al unpubiblished) | NcoI/AflII cut amplicon (primer 0769/0770 template K005) into K167 |
| K293 | Vector for yeast transformation in order to modify ARS315 locus and insert the RS sites next to NS+-2 | Gibson assembly of two fragments primers 815and 816 with pM49.2ARS315+-2 as template and primers 817 and 818 with K174 as template |
| K273 | Vector for yeast transformation in order to modify ARS315 locus and insert the RS sites next to NS+-3 | Gibson assembly of two fragments: primers 819and 820with pM49.2ARS315+-3 as template and primers 821 and 822 with K174 as template |
| K207 | | Bar ko plasmid |
| K303 | Vector for yeast transformation in order to delete the Ies6 gene and replace it with histidine | Gibson assembly of four fragments: Primers 1049 and 1050 , 1053 and 1054 with gDNA as template, Primers 1051 and 1052 with K207 plasmid as template, K18 digested by |
