## Supplementary material for "Single molecule MATAC-seq reveals key determinants of DNA replication origin efficiency": Table of Primers

| | |
|---|---|
| #0016 | GATAAGCTTttttgggtcctttgttttcg |
| #0017 | GATGAATTCatttggagggaggagaagga |
| #0020 | TCAGAATTCtcctccagtgggattgctac |
| #0021 | TCACTGCAGgcctagcgggctattacctt |
| #0040 | ttttgggtcctttgttttcgttgtttcagtctggataaattttaagttac tcgatccgatgataagctgtc |
| #0041 | ATTTGGAGGGAGGAGAAGGATAACAGCGACGAAACACCGGACAGATTCCCtgccacctgacgt ctaagaa |
| #0046 | GAAAGTAGTTATTACGGCGTCGG |
| #0047 | GGCTCTAGGGTAGTTGCG |
| #0048 | gcgacgcccgacgccgtaataactactttcTCGACCCGAGATCATATC |
| #0049 | caatgagagaaacgcaactaccctagagccCCACGATTTGATGAAAGAATAAC |
| #0050 | GTGTGCTAAGTGTCCTGTTTC |
| #0051 | AATATTGTCTTTGGACGTTTG |
| #0052 | gaacgttccgaaacaggacacttagcacacTCGACCCGAGATCATATC |
| #0053 | tggtttgggcaaacgtccaaagacaatattCCACGATTTGATGAAAGAATAAC |
| #0074 | GAAAAGGGCATGTAATATTG |
| #0075 | CTTGGGGAAGAAGTAACAATGAC |
| #0080 | tcctccagtgggattgctacttcttttgttgctgctgcatcctcaacttg tcgatccgatgataagctgtc |
| #0081 | gcctagcgggctattaccttgtaaataccacactatcaatccttaaatgt tgccacctgacgtctaagaa |
| #0119 | aaatcgtggtcgaccggcatgcaagctcccTCGAGGACAGACCACTTATG |
| #0120 | aaccagtggttatatgtacagtagtacgttATCTCTCCGCCTGAATAAG |
| #0121 | aaatcgtggtcgaccggcatgcaagctcccTCGAGAAATACAGAATAGGAAAG |
| #0122 | aaccagtggttatatgtacagtagtacgttATCGTAGCGGTGTTTATC |
| #0223 | Ggcttttcgatcagacttggcatgtgactaatcaagtatggcatgctggt ttttgggtcctttgttttcg |
| #0224 | Tagtaaataacggagactggcgaaccgaatgggcacctgcctctgactgc atttggagggaggagaagga |
| #0251 | TAACTTCAGCACCAAAGCCAACAACTACGACCTATGTCGAGCAACGACTTTCCTCCAGT GGGATTGCTAC |
| #0252 | TTCTTGGCAGTCACATATATGGAAGGTGAATTTAGAGTAGTTTCCTTATAGCCTAGCGGG CTATTACCTT |
| #0270 | TCAgctagcttaattaaAAGACAACAGATTTATTGTA |
| #0271 | TCAggcgcgccCCCGAGGATTATAATTGTTC |
| #0272 | tcaGGCGCGCCtttcgtctcgcgcgtttcgg |
| #0273 | tcaGCTAGCagtagttggaatatcataat |
| #0274 | tcaGCTAGCgagaatttgtattttcaggg |
| #0275 | tcaTTAATTAAccccgttccacaacacaaca |
| #0319 | AAC GAATTC aaaaaccccagcagcagata |
| #0320 | AAC AAGCTT ctcctgggccttgatgatac |
| #0321 | gtggtcgaccggcatgcaagctccctttcctgggcggtttggttac |
| #0322 | gtggttatatgtacagtagtacgttctaactgtaggcgcttttatctcc |
| #0323 | AAC GAATTC GGGAAGGAGATCTGCAGTGT |
| #0324 | AAC AAGCTT CGATTGAAGCCGTGGATGAG |
| #0330 | gtggttatatgtacagtagtacgttGGTTGGGGTATTAAGAGTAC |
| #0331 | gtggtcgaccggcatgcaagctcccTTTTGTATGAAAAACTCATGAATC |
| #0332 | gtggttatatgtacagtagtacgttCTTGGTGCAAGAAGTTGG |
| #0333 | gtggtcgaccggcatgcaagctcccGCCATGCCTAAGAAAGATTG |
| #0334 | gtggttatatgtacagtagtacgttAGGCAGTTTCATCTTCAG |
| #0338 | Agaaagtgcttttggatcgtccggtgaaattgcagtaataccgatagtcc tcgatccgatgataagctgtc |
| #0346 | gaccgataaaaagggtaatagatggCCACGATTTGATGAAAGAATAAC |
| #0347 | ttgatactacaggagctggagatacTCGACCCGAGATCATATC |
| #0350 | tcagcaatcatcattattccCCACGATTTGATGAAAGAATAAC |
| #0351 | atatcgtaaaatttgtgagcTCGACCCGAGATCATATC |

| | |
|---|---|
| #0366 | gggtggtatagcccttggtaCCACGATTTGATGAAAGAATAAC |
| #0367 | ataacccctcacctttcaagTCGACCCGAGATCATATC |
| #0368 | CTTGAAAGGTGAGGGGTTATATAC |
| #0369 | TACCAAGGGCTATACCAC |
| #0457 | atactaattgaagagaaagctggtggccaaaataggatattgattgtaga tgccacctgacgtctaagaa |
| #0458 | Gagcttttctttcctctctctttttttttcttgttacatattcctatat tgccacctgacgtctaagaa |
| #0769 | tacgccaacttaagaccatg |
| #0770 | tctcttttccatggtcatga |
| #0775 | gtggtcgaccggcatgcaagctcccaaaaaaccgcgaagagctcc |
| #0776 | gtggttatatgtacagtagtacgttctcaaccgcagaacccgg |
| #0777 | gtggtcgaccggcatgcaagctcccggttggggtattaagagtacaatgc |
| #0778 | gtggttatatgtacagtagtacgttcgcaattttcttgaacggttttttc |
| #0815 | aaaaaatccggaacaaaaaaaccgcTCGACCCGAGATCATATC |
| #0816 | ACCAACTAATTACTGCTCAACCGCACCACGATTTGATGAAAGAATAAC |
| #0817 | acgttattctttcatcaaatcgtggTGCGGTTGAGCAGTAATTAG |
| #0818 | ccacagtgatatgatctcgggtcgaGCGGTTTTTTTGTTCCGG |
| #0819 | actctcactattttgtttttcgacccgagatcatatc |
| #0820 | cgtatatgactgcacaagacccacgatttgatgaaagaataac |
| #0821 | gtcttgtgcagtcatatac |
| #0822 | aaaaacaaaatagtgagagtaatg |
| #0848 | AATCTCACTAAAAGTAACATACAGTACCGATAAATCGAGATTGCAGAGTA tgccacctgacgtctaagaa |
| #0849 | GTTCAATTATCTTAGAATGGATATGAATTAGTTAAAGCGGCTCGACCCAG tcgatccgatgataagctgtc |
| #0858 | TCCATGTCCATGTCCATGTCATCATGGGCCGTGACAAGCGTCGCCGCGCA gccgaataaacttaaaattga |
| #0859 | CCTCGACGGCCTCCAGTTCTTCGACCAACTGTTCGTGATCGTCATCCATT gagcttttctttcctctctct |
| #0932 | attcaaatacgtcatcctat |
| #0933 | actggatacttggtcgtcgt |
| #1049 | gggtaccgggccccccctcgaggtcgacggtatcgataagcttgatatcgcctgcaggagaaaaaaagaatagtaataacaatttg |
| #1050 | acattatatgacccttctagacacttgtttcgactttctcacg |
| #1051 | tcatcagcgtgagaaagtcgaaacaagtgtctagaagggtcatataatg |
| #1052 | aaatacatacatacatatacaatgcaatcaaaattgtattccacgtc |
| #1053 | ggtgacgtggaatacaattttgattgcattgtatatgtatgtatgtattttag |
| #1054 | ccaccgcggtggcggccgctctagaactagtggatcccccgggctgcagggagctcgttctcccaatgaatggtc |
| #1158 | ccctacaccacagcatacaaaaccgaatcaaaattgtattccacgtc |
| #1159 | ggtgacgtggaatacaattttgattcggttttgtatgctgtgg |
