## Supplementary material for "Single molecule MATAC-seq reveals key determinants of DNA replication origin efficiency": Table of Yeast strains

| | | | |
|---|---|---|---|
| Y0001 (Y01408) | MATa; ura3Δ0; leu2Δ0; his3Δ1; met15Δ0; bar1::kanMX4 | Wildtype strain | EUROSCARF |
| Y0008 (ySH089) | MATa; ade2-1; ura3-1; trp1-1; leu2-3,112; his3-11; can1-100 | URA3::LEU2 pTEF2 LEXA-TAP pGAL RecR | Transformation of SbfI digested fragment of pSH21 for genomic integration of LEU2 pTEF2 LexA-TAP pGAL RecR expression cassette |
| Y0010 (yMW2) | MATa; ura3Δ0; leu2Δ0; his3Δ1; met15Δ0; bar1::kanMX4; ARS305::URA3 | ARS305 exchanged for URA3 | Transformation of amplicon derived from K001 (primer 0040/0041) into Y0001. Selection on Ura |
| Y0011 (yMW3) | MATa; ura3Δ0; leu2Δ0; his3Δ1; met15Δ0; bar1::kanMX4; ARS316::URA3 | ARS316 exchanged for URA3 | Transformation of amplicon derived from K001 (primer 0080/0081) into Y0001. Selection on Ura |
| Y0018 (yMW6) | MATa; ura3Δ0; leu2Δ0; his3Δ1; met15Δ0; bar1::kanMX4; RS_LEXA_NS-3_ARS305_NS+3_RS | RS sites and lexA binding sites at ARS305 after +-3 nucleosomes | Transformation of amplicon derived from K113 (primer 0223/0224) into Y0010. Selection on FOA |
| Y0021 (yMW9) | MATa; ura3Δ0; leu2Δ0; his3Δ1; met15Δ0; bar1::kanMX4; RS_LEXA_NS-3_ARS316_NS+3_RS | RS sites and lexA binding sites at ARS316 after +-3 nucleosomes | Transformation of amplicon derived from K116 (primer 0251/0252) into Y0011. Selection on FOA |
| Y0042 (yAC02) | MATa; ura3Δ0; leu2Δ0; his3Δ1; met15Δ0; bar1::kanMX4; ARS313::URA3 | ARS313 exchanged for URA3 | Transformation of amplicon derived from K001 (primer 0338/0457) into Y0001. Selection on Ura |
| Y0043 (yAC03) | MATa; ura3Δ0; leu2Δ0; his3Δ1; met15Δ0; bar1::kanMX4; ARS315::URA3 | ARS315 exchanged for URA3 | Transformation of amplicon derived from K001 (primer 0340/0458) into Y0001. Selection on Ura |
| Y0044 (yAC04) | MATa; ura3Δ0; leu2Δ0; his3Δ1; met15Δ0; bar1::kanMX4;RS_LEXA_NS-3_ARS313_NS+3_RS | RS sites and lexA binding sites at ARS313 after +-3 nucleosomes | Transformation of EcoRI/HindIII digested plasmid K170 into Y0042. Selection on FOA |
| Y0065 (yTS3) | MATa; ura3Δ0; leu2Δ0; his3Δ1; met15Δ0; bar1::kanMX4; RS_LEXA_NS-3_ARS305_NS+3_RS; Chr I 212kb::LEU2 pTEF2-LEXA-TAP pGAL1-10 RecR | RS sites and lexA binding sites at ARS305 after +-3 nucleosomes. Expression cassette for R Recombinase and lexA (TEF2 promoter) | Transformation of SbfI digested plasmid K238 into Y0018. Selection on Leu |

| | | | |
|---|---|---|---|
| Y0066 (yTS4) | MATa; ura3Δ0; leu2Δ0; his3Δ1; met15Δ0; bar1::kanMX4; Chr I 212kb::LEU2 pTEF2-LEXA-TAP pGAL1-10 RecR | WT strain with expression cassette for R Recombinase and lexA (TEF2 promoter) | Transformation of SbfI digested plasmid K238 into Y0001. Selection on Leu |
| Y0069 (yMW38) | MATa; ura3Δ0; leu2Δ0; his3Δ1; met15Δ0; bar1::kanMX4; RS_LEXA_NS-3_ARS316_NS+3_RS; Chr I 212kb::LEU2 pTEF2-LEXA-TAP pGAL1-10 RecR | RS sites and lexA binding sites at ARS316 after +-3 nucleosomes. Expression cassette for R Recombinase and lexA (TEF2 promoter) | Transformation of SbfI digested plasmid K238 into Y0021. Selection on Leu |
| Y0084 (yMW53) | MATa; ade2-1; ura3-1; trp1-1; leu2-3,112; his3-11; can1-100; bar1::his3 | Bar1 knockout | Transformation of amplicon derived from K207 (primers 932/933) into Y8. Selection on His |
| Y0089 (yAC14) | MATa; ura3Δ0; leu2Δ0; his3Δ1; met15Δ0; bar1::kanMX4;RS_LEXA_NS-3_ARS315_NS+3_RS | RS sites and lexA binding sites at ARS315 after +-3 nucleosomes | Transformation of amplicon derived from K273 (primer 0858/0859) into Y0043. Selection on FOA |
| Y0091 (yAC16) | MATa; ura3Δ0; leu2Δ0; his3Δ1; met15Δ0; bar1::kanMX4;RS_LEXA_NS-3_ARS315_NS+3_RS; Chr I 212kb::LEU2 pTEF2-LEXA-TAP pGAL1-10 RecR | RS sites and lexA binding sites at ARS315 after +-3 nucleosomes. Expression cassette for R Recombinase and lexA (TEF2 promoter) | Transformation of SbfI digested plasmid K238 into Y0089. Selection on Leu |
| Y0094 (yAC19) | MATa; ura3Δ0; leu2Δ0; his3Δ1; met15Δ0; bar1::kanMX4;RS_LEXA_NS-3_ARS313_NS+3_RS; Chr I 212kb::LEU2 pTEF2-LEXA-TAP pGAL1-10 RecR | RS sites and lexA binding sites at ARS313 after +-3 nucleosomes. Expression cassette for R Recombinase and lexA (TEF2 promoter) | Transformation of SbfI digested plasmid K238 into Y0044. Selection on Leu |
| Y0102 (AC20) | MATa; ura3Δ0; leu2Δ0; his3Δ1; met15Δ0; bar1::kanMX4;RS_LEXA_NS-3_ARS315_NS+3_RS; Chr I 212kb::LEU2 pTEF2-LEXA-TAP pGAL1-10 RecR ; isw2Δ::ura3 | Deletion of isw2 gene | Transformation of amplicon derived from K1 (primers 848/849) into Y0091. Selection on URA |
| Y0103 (AC21) | MATa; ura3Δ0; leu2Δ0; his3Δ1; met15Δ0; bar1::kanMX4;isw2Δ::ura3 | Deletion of isw2 gene | Transformation of amplicon derived from K1 (primers 848/849) into Y0001. Selection on URA |
| Y0104 (AC22) | MATa; ura3Δ0; leu2Δ0; his3Δ1; met15Δ0; bar1::kanMX4; Chr I 212kb::LEU2 pTEF2-LEXA-TAP pGAL1-10 RecR ;isw2Δ::ura3 | Deletion of isw2 gene | Transformation of amplicon derived from K1 (primers 848/849) into Y0066. Selection on URA |
| Y0105 (AC23) | MATa; ura3Δ0; leu2Δ0; his3Δ1; met15Δ0; bar1::kanMX4; | Deletion of isw2 gene | Transformation of amplicon derived from K1 (primers 848/849) |

| | | | |
|---|---|---|---|
| | RS_LEXA_NS-3_ARS305_NS+3_RS; Chr I 212kb::LEU2 pTEF2-LEXA-TAP pGAL1-10 RecR ;isw2Δ::ura3 | | into Y0065. Selection on URA |
| Y0106 (AC24) | MATa; ura3Δ0; leu2Δ0; his3Δ1; met15Δ0; bar1::kanMX4; RS_LEXA_NS-3_ARS316_NS+3_RS; Chr I 212kb::LEU2 pTEF2-LEXA-TAP pGAL1-10 RecR;isw2Δ::ura3 | Deletion of isw2 gene | Transformation of amplicon derived from K1 (primers 848/849) into Y0069. Selection on URA |
| Y0107 (AC25) | MATa; ura3Δ0; leu2Δ0; his3Δ1; met15Δ0; bar1::kanMX4;RS_LEXA_NS-3_ARS313_NS+3_RS; Chr I 212kb::LEU2 pTEF2-LEXA-TAP pGAL1-10 RecR;isw2Δ::ura3 | Deletion of isw2 gene | Transformation of amplicon derived from K1 (primers 848/849) into Y0069. Selection on URA |
| Y127 (AC37) | MATa; ura3Δ0; leu2Δ0; his3Δ1; met15Δ0; bar1::kanMX4; RS_LEXA_NS-3_ARS305_NS+3_RS; Chr I 212kb::LEU2 pTEF2-LEXA-TAP pGAL1-10 RecR ;ies6Δ::his | Deletion of ies6 gene | Transformation of SbfI and SacI digested plasmid K303 into Y0065. Selection on histidine |
| Y128 (AC38) | MATa; ura3Δ0; leu2Δ0; his3Δ1; met15Δ0; bar1::kanMX4;RS_LEXA_NS-3_ARS313_NS+3_RS; Chr I 212kb::LEU2 pTEF2-LEXA-TAP pGAL1-10 RecR ;ies6Δ::his | Deletion of ies6 gene | Transformation of SbfI and SacI digested plasmid K303 into Y0094. Selection on histidine |
| Y129 (AC39) | MATa; ura3Δ0; leu2Δ0; his3Δ1; met15Δ0; bar1::kanMX4; RS_LEXA_NS-3_ARS316_NS+3_RS; Chr I 212kb::LEU2 pTEF2-LEXA-TAP pGAL1-10 RecR;ies6Δ::his | Deletion of ies6 gene | Transformation of SbfI and SacI digested plasmid K303 into Y0069. Selection on histidine |
| Y130 (AC40) | MATa; ura3Δ0; leu2Δ0; his3Δ1; met15Δ0; bar1::kanMX4;RS_LEXA_NS-3_ARS315_NS+3_RS; Chr I 212kb::LEU2 pTEF2-LEXA-TAP pGAL1-10 RecR ;ies6Δ::his | Deletion of ies6 gene | Transformation of SbfI and SacI digested plasmid K303 into Y0091. Selection on histidine |
| Y135 (AC44) | MATa; ura3Δ0; leu2Δ0; his3Δ1; met15Δ0; bar1::kanMX4; Chr I 212kb::LEU2 pTEF2-LEXA- | Deletion of ies6 gene | Transformation of SbfI and SacI digested plasmid K303 into Y0066. Selection on histidine |

| | | | |
|---|---|---|---|
| | TAP pGAL1-10 RecR ;ies6Δ::his | | |
| Y136 (AC45) | MATa; ade2-1; ura3-1; trp1-1; leu2-3,112; his3-11; can1-100; bar1::his3; isw2Δ::ura3 | Deletion of isw2 gene | Transformation of amplicon derived from K1 (primers 848/849) into Y0084. Selection on URA |
